# Coevolution of Patch Selection in Stochastic Environments

**DOI:** 10.1101/2022.03.14.484312

**Authors:** Sebastian J. Schreiber, Alexandru Hening, Dang H. Nguyen

## Abstract

Species interact in landscapes where environmental conditions vary in time and space. This variability impacts how species select habitat patches. Under equilibrium conditions, coevolution of this patch selection can result in ideal-free distributions where per-capita growth rates are zero in occupied patches and negative in unoccupied patches. These ideal-free distributions, however, don’t explain why species occupy sink patches, competitors have overlapping spatial ranges, or why predators avoid highly productive patches. To understand these patterns, we analyze multi-species Lotka-Volterra models accounting for spatial heterogeneity and environmental stochasticity. In occupied patches at the coESS, we show that the differences between the local contributions to the mean and the variance of the long-term population growth rate are equalized. Applying this characterization to models of antagonistic interactions reveals that environmental stochasticity can partially exorcize the ghost of competition past, select for new forms of enemy-free and victimless space, and generate Hydra effects over evolutionary time scales. Viewing our results through the economic lens of Modern Portfolio Theory highlights why the coESS for patch selection is often a bet-hedging strategy coupling stochastic sink populations. Our results highlight how environmental stochasticity can reverse or amplify evolutionary outcomes due to species interactions or spatial heterogeneity.

## Introduction

Evolution of habitat choice plays a key role in shaping the distribution and abundance of species. Evolutionary drivers of this choice include spatial and temporal variation in abiotic conditions among habitat patches, and species interactions within habitat patches. One successful theoretical approach to evaluate the relative importance of these drivers assumes that individuals can freely chose their habitat patches with no costs to dispersal (Fretwell and Lucas, 1969; Rosenzweig, 1981, 1991; Holt, 1997; Morris, 2003; Křivan et al., 2008; Morris, 2011). Under equilibrium conditions, this theoretical approach provides empirically supported predictions about the spatial distributions of predators and their prey (Oksanen et al., 1995; Schreiber et al., 2000) and competing species (Lawlor and Maynard Smith, 1976; Diamond, 1978; Connell, 1980). In temporally variable environments, this approach also provides an evolutionary explanation of why populations occupy habitat patches where deaths exceed births (Holt, 1997; Jansen and Yoshimura, 1998; Schreiber, 2012). Despite these significant advances, there isn’t a comprehensive approach for how temporal variation, spatial heterogeneity, and species interactions simultaneously drive the coevolution of habitat choices. Here, we introduce one such framework.

When individuals select habitat patches to maximize their fitness, theory predicts that the population will reach an ideal-free distribution in which the per-capita growth rates are equal in all of the occupied patches and lower in the unoccupied patches (Fretwell and Lucas, 1969; Křivan et al., 2008). Two classical concepts, enemy-free space and the ghost of competition past, from evolutionary ecology follow from this ideal-free theory. Jeffries and Lawton (1984) defined enemy-free space as “ways of living that reduce or eliminate a species’ vulnerability to one or more species of natural enemies.” In a spatial context, enemy-free space corresponds to a species living in habitat patches where there are fewer or no natural enemies, a phenomena that has been observed in several empirical systems (Denno et al., 1990; Fox and Eisenbach, 1992; Berdegue et al., 1996; Murphy, 2004; Cole et al., 2005; Heisswolf et al., 2005; Kaminski et al., 2010; Roy et al., 2011; Murphy et al., 2014; Greeney et al., 2015). Ideal-free distributions of predators and their prey yield enemy-free space when patches of lower quality for the prey are also lower quality for the predator (Schreiber et al., 2000; Schreiber and Vejdani, 2006). Consistent with these theoretical predictions, Fox and Eisenbach (1992) found that diamondback moth *Plutella xylostella* preferentially laid eggs on collards and red cabbage grown on low-fertilized soils while its main parasitoid, an ichneumonid wasp *Diadegma insulare*, preferentially searched for hosts on collards grown on high-fertilized soils. These contrary choices occurred despite diamond back moth larvae, in the absence of the wasps, having higher survival rates and growing to larger sizes on host plants from high-fertilized soils.

For competing species, ideal-free theory predicts that, eventually, competitors never occupy the same habitat patch. At equilibrium, each species only occupies patches in which they are competitively superior (Lawlor and Maynard Smith, 1976). As competitive interactions from the past led to habitat choices eliminating competition in the present, this outcome is known as “the ghost of competition past” (Connell, 1980). Such a haunting may explain the spatial distribution of two *Crateroscelis* warblers species in New Guinea (Diamond, 1973, 1978) where one species abruptly replaces the other at an altitude of 1, 643 meters. Despite this sharp transition in warblers, shifts in abundance of competing species typically are more gradual with substantive regions of overlap (Noon, 1981; Chettri and Acharya, 2010; Campos-Cerqueira et al., 2017; Burner et al., 2019). Along these regions of overlap, each competitor may shift from living in source patches (e.g. where it is competitively dominant) to living in sink patches (e.g. where it is competitively inferior) (Amarasekare and Nisbet, 2001).

Under equilibrium conditions, ideal-free theory predicts birth rates equal death rates in occupied patches. Consequently, there can be no sink populations – local populations whose death rates exceed their birth rates (Holt, 1985; Pulliam, 1988; Pulliam and Danielson, 1991). None the less, sink populations have been observed in many taxonomic groups including birds (Dias et al., 1996; Vierling, 2000; Keagy et al., 2005; Tittler et al., 2006), plants (Kadmon and Tielbörger, 1999) mammals (Kreuzer and Huntly, 2003; Robinson et al., 2008; Monson et al., 2011), reptiles (Manier and Arnold, 2005), amphibians (Rowe et al., 2001) and fishes (Hänfling and Weetman, 2006; Barson et al., 2009; McDowall, 2010). Indeed, a meta-analysis of 90 source-sink assessments found that 60% of the studied populations were identified as sink populations (Furrer and Pasinelli, 2016). These sink populations can be either unconditional or conditional sink populations (Loreau et al., 2013). Unconditional sink populations have negative per-capita growth rates in the absence of conspecific and antagonistic interactions. Alternatively, conditional sink populations have negative per-capita growth rates due to high densities of conspecifics (Watkinson and Sutherland, 1995), heterospecific competitors (Amarasekare and Nisbet, 2001) or predators (Holt, 1977).

Patch selection theory for single species models suggest that unconditional sink populations may evolve in temporally variable environments (Holt, 1997; Jansen and Yoshimura, 1998; Schmidt et al., 2000; Jonzén et al., 2004; Schreiber, 2012). Intuitively, making use of low quality, but environmentally stable patches can buffer populations against environmental fluctuations in patches that, on average, are of higher quality. (Cohen, 1966; Holt, 1997; Jansen and Yoshimura, 1998; Kisdi, 2002; Schreiber, 2012). However, whether this evolutionary explanation also extends to sink populations in a community context remains largely unexplored. A notable exception is the work of Schmidt et al. (2000) who studied the evolution of patch choice for two competing species in a fluctuating environment with two habitat patches. Schmidt et al. (2000) found that these environmental fluctuations can result in patches being occupied by both competitors and, thereby, partially exorcize the ghost of competition past. However, to what extent these conclusions apply to more than two competing species, or other forms of species interactions, or landscapes of greater complexity is unknown.

Here, we introduce a framework for analyzing the coevolution of patch-selection for multispecies communities in spatially and temporally heterogeneous environments. The framework involves stochastic counterparts of the generalized Lotka–Volterra models that have been a mainstay of theoretical work in community ecology (May, 1975; Holt, 1977; Polis and Holt, 1992; Law and Morton, 1996; Chesson and Kuang, 2008; Edwards and Schreiber, 2010; Rohr et al., 2016; Schreiber et al., 2018). For these models, we explore coevolutionary stable strategies (coESS) for patch-selection whereby each species has its own patch-selection strategy and any subpopulation playing a different strategy fails to establish (Roughgarden, 1979; Brown and Vincent, 1987; Foster and Young, 1990; Rand et al., 1994; Feng et al., 2022). Using an analytically tractable characterization of the coESSs, we examine several questions. First, for species playing the coESS for patch-selection, we ask is there any demographic quantity which is equal in all occupied patches? We provide a positive answer to this question and, therby, extend earlier work on single species models (Schreiber, 2012; Evans et al., 2015). Second, when do sink population evolve? In particular, as earlier theory only considered single species models (Holt, 1997; Jansen and Yoshimura, 1998; Schreiber, 2012; Evans et al., 2015), when do species interactions result in the evolution of conditional versus unconditional sink populations? Finally, in what ways does environmental stochasticity exorcize the ghost of competition past, and what ways does it amplify or dampen the evolution of enemy-free space? Collectively, the answers to these questions highlight the interactive effects of spatial heterogenity, temporal variation, and species interactions on the evolution of habitat choice.

## Models and Methods

We model a community of *n* species living in an environment with *k* patches. These patches may represent distinct habitats, patches of the same habitat type, or combinations thereof. The dynamics within a patch are modeled by stochastic Lotka–Volterra differential equations (May, 1973, 1975; Turelli, 1977, 1978; Turelli and Gillespie, 1980; Turelli, 1986; Lande et al., 2003; Schreiber et al., 2011; Evans et al., 2013, 2015; Nolting and Abbott, 2016; Hening and Nguyen, 2018*c**,b*; Hening et al., 2021). To couple these local dynamics, we do not explicitly model movement but, instead, we assume that each species has a fixed fraction of its population in each of these patches. We call this fixed spatial distribution for a species, its patch-selection strategy. This strategy may correspond to freely dispersing individuals spending a fixed fraction of time in each patch, or allocating a fixed fraction of their offspring to patch (Holt, 1997; Jansen and Yoshimura, 1998; Schmidt et al., 2000; Bascompte et al., 2002; Schreiber, 2012; Evans et al., 2015). The resulting models are a multispecies version of the models introduced in (Schreiber, 2012).

We introduce a definition of coevolutionary stable strategies (coESS) of patch-selection. Our definition merges the concepts of coESS for deterministic models (Roughgarden, 1979; Brown and Vincent, 1987; Rand et al., 1994) and ESS for single species stochastic models (Foster and Young, 1990; Feng et al., 2022). Roughly, coevolutionary stability requires that any small population of a species playing a different strategy from the rest of its popualtion will not establish. Using invasion growth rates of mutant strategies, we provide both analytical and numerical approaches for computing the coESS.

### The Models

#### The within-patch dynamics

Consider one patch in the landscape, say patch *ℓ*. For the moment, assume that all species only use this patch i.e. the patch-selection strategy of each species *i* is to have 100% of individuals use patch *ℓ*. To model the dynamics of species *i* within this patch, let *x*_*i*_(*t*) denote its density at time *t*, 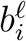 its intrinsic per-capita growth rate in the absence of other species, and 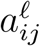 its per-capita interaction rate with species *j*. These quantities determine the deterministic forces acting on species *i*. Specifically, the change Δ*x*_*i*_(*t*) = *x*_*i*_(*t* + Δ*t*) − *x*_*i*_(*t*) in the density of species *i* over a small time step Δ*t* satisfies

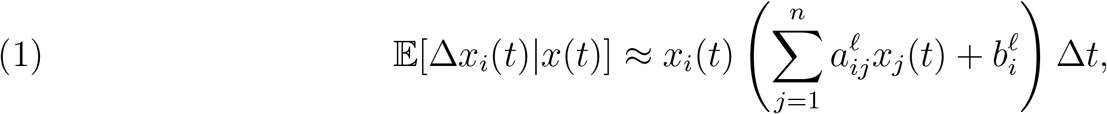

where *x* = (*x*_1_(*t*), …, *x*_*n*_(*t*)) is the community state in patch *ℓ* at time *t*, and 𝔼[*X* |*Y*] denotes the conditional expectation of a random variable *X* with respect to the random variable *Y*. Thus, the expected instantaneous change of the species density is given by a Lotka-Volterra model.

To capture the role of environmental stochasticity, we assume that the variance in the growth of species *i* in patch *ℓ* over a time interval Δ*t* satisfies

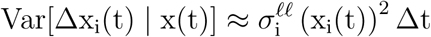

where Var[*X* |*Y*] denotes the conditional variance of *X* with respect to *Y*. Taking the limit as Δ*t* gets infinitesimally small, the population dynamics when all individuals only use patch *ℓ* are given by the Itô stochastic differential equations (Gardiner, 2009; Oksendal, 2013)

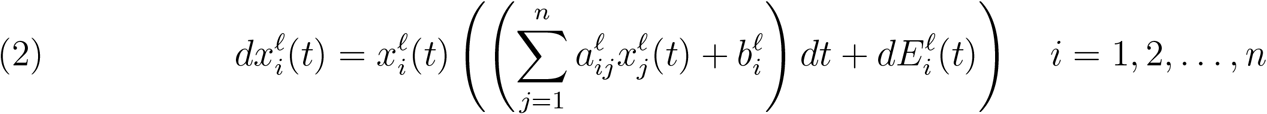

where 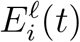 is a (non-standard) Brownian motion with mean 0 and variance 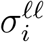 (see Appendix A for how to represent *E*_*i*_(*t*) as a linear combination of standard Brownian motions). One can interpret (2) as approximately updating densities by Lotka-Volterra dynamics plus multiplicative, normally distributed noise that corresponds to fluctuations in the intrinsic rates of growth.

#### The global dynamics

To describe the global dynamics when species use more than one patch, let 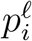 denote the fraction of individuals of species *i* selecting patch *ℓ* and 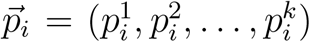 denote the patch selection strategy of species *i* e.g. if 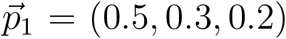 then there are *k* = 3 patches and, at any point in time, 50% of individuals of species 1 are using patch 1, 30% are using patch 2, and 20% are using patch 3. If *x*_*i*_ denotes the global density of species *i*, then 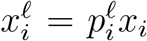 is the density of species *i* in patch *ℓ*. Let 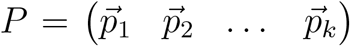 denote the matrix of the patch selection strategies for all species where the *i*-th column of *P* corresponds to the patch selection strategy of species *i*.

To account for spatial correlations in the environmental fluctuations across the patches, we assume that the per-capita growth rates of the species *i* in patches *ℓ* and *m* over a time interval of length Δ*t* satisfy

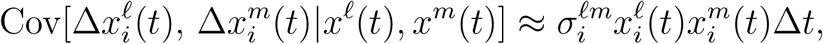

where Cov[*X, Y*|*Z, W*] denotes the covariance between random variables *X* and *Y* given the random variables *Z* and *W*. The covariance matrix 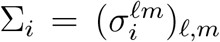 for species *i* captures the spatial dependence between the temporal fluctuations in intrinsic growth rates across patches.

Under these assumptions, the community dynamics of the *n* species interacting in the *k* patches are given by the system of Itô stochastic differential equations

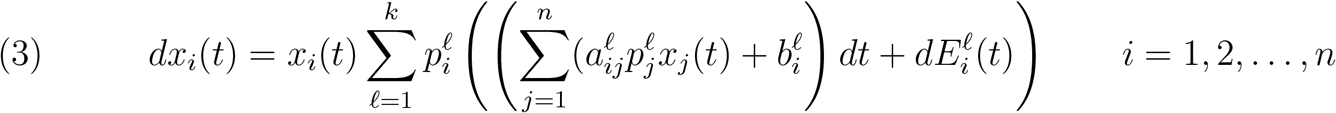

where for species *i*, 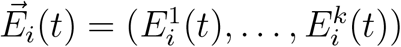 is a multivariate Brownian motion with covariance matrix Σ_*i*_. We make no assumptions about the cross-correlations, 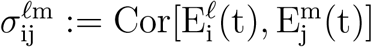, in the environmental fluctuations experienced by different species *i* ≠ *j*.

### Methods

To study coevolution of patch-selection strategies, we introduce several methods. First, we characterize the mean densities of the species at stationary distributions for the community. Second, we introduce a definition of a coevolutionary stable strategy that acounts for environmental stochasticity. Finally, we describe a numerical method for solving for these coevolutionarily stable strategies.

#### Stationary distributions and stochastic growth rates

In order to study the coevolution of patch-selection strategies, we need to identify when species coexist. Criteria for determining coexistence for stochastic Lotka-Volterra models were developed by Hening and Nguyen (2018*a*); Hening et al. (2021), complementing earlier work for discrete-time models and continuous-time replicator equations (Schreiber et al., 2011; Benaïm and Schreiber, 2019). These criteria are based on invasion growth rates, the long-term average growth rates of species when rare (see Appendix A). Importantly, these invasion based criteria ensure that the statistical properties of the coexisting species’ densities are characterized by a unique stationary distribution (depending on *P*). When the patch-selection strategies of the species are 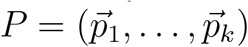, let 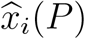 be the mean density of species *i* at this stationary distribution, and 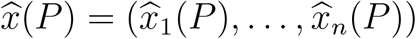.

At this stationary distribution, *the local long-term growth rate of species i in patch ℓ* equals the difference between its average, local per-capita growth rate and one-half of the environmental variance experienced in patch *ℓ*:

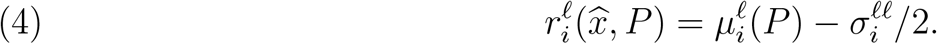

where

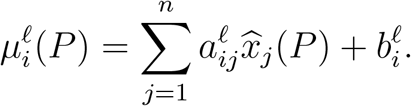

This local long-term growth rate describes the per-capita growth of a population averaged across the fluctuations in species’ densities and environmental conditions. As the per-capita impacts of the environmental fluctuations on the per-capita growth rates are density-independent, the reduction 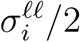 in the local, long-term growth rates are also density-independent.

This local long-term growth rate characterizes the rate of growth for a subpopulation of species *i* permanently restricted to patch *ℓ*. If this local long-term growth rate is negative, such a subpopulation of such individuals would exponentially decline to extinction. Whenever this occurs, patch *ℓ* is a long-term sink for species *i*. When the local long-term growth rate is negative due to the environmental fluctuations (i.e. 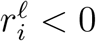 despite 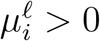), growth rates in these patches fluctuate between positive and negative values over shorter time scales. Hence, we call these long-term sinks, stochastic sinks. Alternatively, if this local long-term growth rate is positive, then this patch is a long-term source patch for species *i*. As we discuss in the results, all patches may be sink patches for species *i* despite species *i* persisting.

At the global scale, the global long-term growth rate of any of the coexisting species must equal zero as their average densities in the long-term are not changing. As this global long-term growth rate is given by the difference between the mean growth rate *M*_*i*_(*P*) and one-half of the global environmental variance *V*_*i*_(*P*) experienced by species *i*, we have

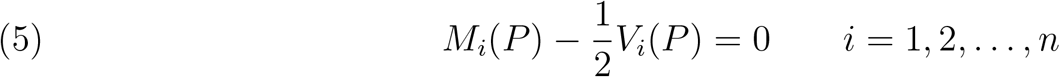

where

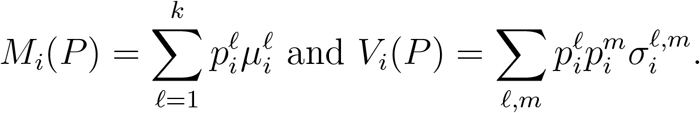

Importantly, equation (5) is a system of linear equations that allows one to easily solve for the mean species densities 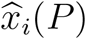. See Appendix A for a proof of (5).

##### Coevolutionary Stable Strategies

To define a coevolutionarily stable strategy *P* of patch-selection, we consider a resident community of coexisting species playing patch-selection strategy *P*. For one of these species, say species *i*′, a mutation arises leading to patch-selection strategy 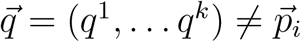. If *y* is the global population density of the mutant and *x*_*i*′_ is the density of the non-mutant individuals of species *i*′, then the resident-mutant community dynamics become

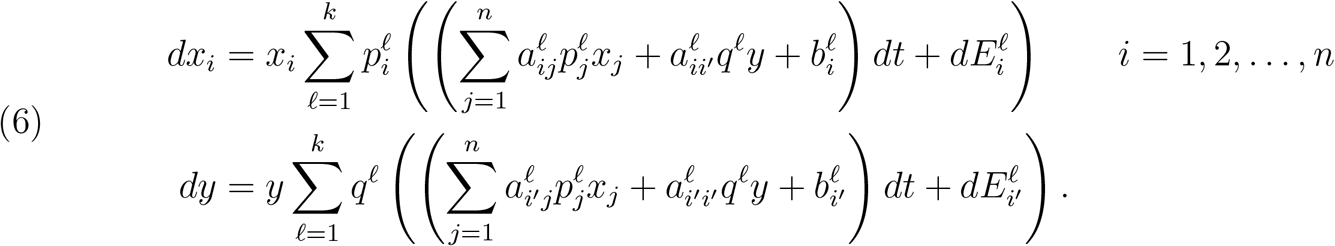

Importantly, the mutant *y* only differs from the resident in its patch-selection strategy 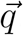. It has the same interaction coefficients *a*_*i*′*j*_ and intrinsic rates of growth *b*_*i*′_ as the resident *x*_*i*′_, and experiences the same environmental stochasticity 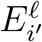 as the resident *x*_*i*′_. When mutant population density is low and the resident community coexist about a stationary distribution, the long-term growth rate, also known as the invasion growth rate, of the mutant population against the resident community is

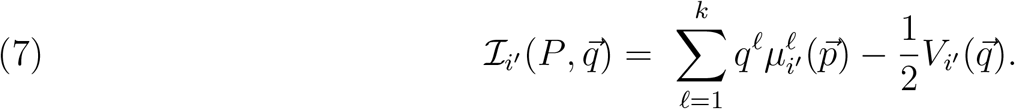

If 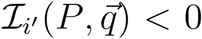, then the mutant is very likely to go asymptotically extinct. In particular, the probability of the mutant going asymptotically extinct is arbitrarily close to one when its initial density is arbitrarily small (see Appendix B).

We define *P* to be a *coevolutionary stable strategy (coESS)* if for every species *i*, mutants playing a different strategy 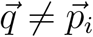 can not invade the community i.e. 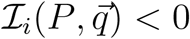 (see Appendix B). When the environmental fluctuations are not perfectly correlated between any pair of patches (i.e. the covariance matrices Σ_*i*_ are non-degenerate), we show in Appendix B that it suffices to check the weaker condition: 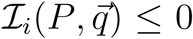 for all *i* and 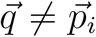. This condition is easier to verify algebraically.

##### Analytical and numerical approaches for the coESS

Our Nash equilibrium condition for the coESS and the explicit analytical expressions for the invasion growth rates can be used in conjunction with the method of Lagrange multipliers to identify a necessary condition for the coESS. This result and its implications are presented in the results section.

To solve for the coESS numerically, we describe in Appendix C an evolutionary dynamic on the strategy space for all of the species in which small mutations occurring at a rate *ν* randomly shuffle the “infinitesimal” weights of our species’ patch selection strategy. This results in a replicator-type equation

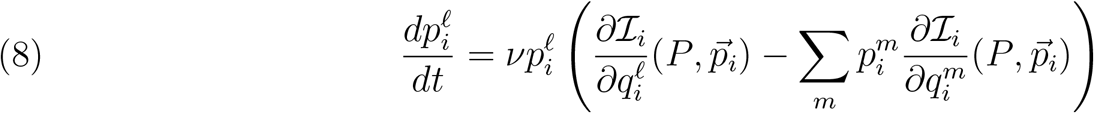

where 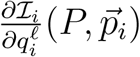 denotes 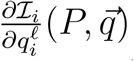 evaluated at 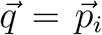. Equation (8) is a multispecies version of the trait dynamics derived in (Schreiber, 2012). In Appendix C, we show that equilibria of (8) satisfy the derivative conditions for a coESS. We simulate (8) using the deSolve package from R (Soetaert et al., 2010). In all cases, these simulations converged to an equilibrium that satisfied the necessary conditions for a coESS. We note that even though the right hand side of (8) for a fixed value of *P* corresponds to the gradient of the function 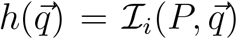 with respect to the Shahshahani metric (Hofbauer and Sigmund, 1998), this is not a gradient ascent method as a coESS (like an ESS) need not maximize the function ℐ_*i*_.

## Results

We begin by presenting a characterization of the coESS and its implications for any number of species. This characterization provides answers to the questions: What, if any, quantities are equalized across the occupied patches? When is there evolution of sink populations? When and how do occupying multiple patches buffer against temporal variability? After answering these questions, we focus on models of antagonistic interactions.

### Characterization of the coESS and General Implications

#### The coESS balances local contributions to the global long-term growth rates

Our analysis reveals that for species playing the coESS, there is a demographic quantities associated with each patch that is equalized across all occupied patches. This demographic quantity corresponds to the difference between the local contributions to the mean and variances of the global long-term growth rate. Consequently, occupied patches with higher contributions to the mean also experience higher contributions to the variance. Intuitively, if there were a patch that contributed relatively more to the mean than the variance, then a mutant subpopulation allocating more individuals to this patch would simultaneously increase their mean rate of growth and decrease the variance in their growth rate. Thus, this mutant would have a higher global long-term growth rate than the residents and could invade.

To describe these local contributions precisely, recall that the global long-term growth rate of species *i* equals the difference between the global mean 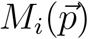 and one-half of the global variance 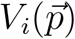. As the mean of the global growth rate is the weighted combination of the local growth rates i.e. 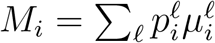, we call 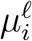 the contribution of patch *ℓ* to *M*_*i*_. The variance means of the of the global growth rate equals the sum of the environmental covariances 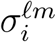 weighted by the probability that two randomly chosen individuals are in patches *ℓ* and *m*:

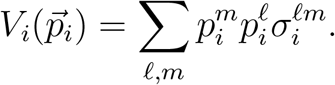

The variance *V*_*i*_ can be expressed as a weighted sum of the covariances between the environmental fluctuations experienced in patch *ℓ* by species *i* and the environmental fluctuations experienced by a randomly chosen individual of species *i*. This covariance for species *i* in patch *ℓ* equals

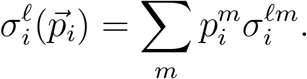

The global variance *V*_*i*_ equals the weighted sum of these covariances

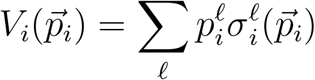

and, therefore, we call 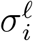 the contribution of patch *ℓ* to *V*_*i*_.

For each species playing the coESS, we show in Appendix D that the difference between the local contributions, 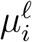 and 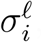, to the mean global growth rate *M*_*i*_ and the global variance *V*_*i*_ are equal in all occupied patches. Moreover, the common value of these differences equals the difference between the global growth rate *M*_*i*_ and the global variance *V*_*i*_. Mathematically,

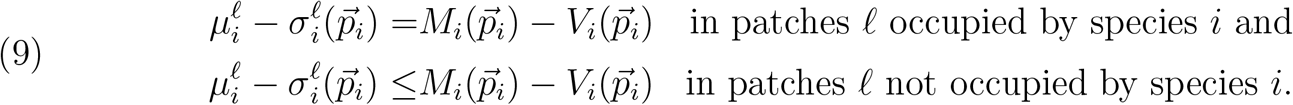

For the unoccupied patches, inequality (9) is strict whenever Σ_*i*_ is non-degenerate, e.g. the environmental fluctuations for species *i* are positive in all patches and no pair of patches has perfectly correlated environmental fluctuations.

Using the coESS conditions (9), we can answer three questions about patch selection: When does the coESS correspond to an ideal-free distribution? When do species evolve to use multiple habitat patches? When is there selection for spatial buffering?

#### Ideal-free distributions are the exception, not the norm

Consistent with classical idealfree distribution theory, when there are no environmental fluctuations (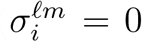 for all *ℓ, m*), the coESS condition (9) implies that the local long-term growth rates are equal in all occupied patches (i.e. 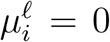 in all occupied patches as 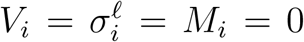) and lower in unoccupied patches (i.e. 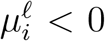). In fluctuating environments, however, the local long-term growth rates, in general, need not be equal in all patches. This occurs as the non-local quantities 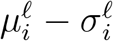 in coESS conditions (9) typically do not equal the local long-term growth rates 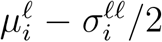. One important exception occurs when the environmental fluctuations are perfectly correlated and have the same magnitude across all patches, i.e. 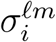 are equal for all *ℓ, m*. In this case, the covariances 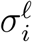, the local variances 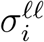, and the global variance *V*_*i*_ are all equal. Consequently, the coESS condition (9) implies that the local long-term growth rates 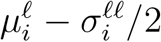 equal zero in all of the occupied patches. Figure 3B-D illustrates how this ideal-free distribution breaks down with reduced spatial correlations in a predator-prey system.

#### Patches acting as environmental buffers are long-term, deterministic sinks

A patch buffers a species against environmental fluctuations when the fluctuations within the patch are negatively correlated with the average fluctuations experienced by the species, i.e. 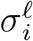 is negative. The coESS condition (9) provides some insights into when such buffering occurs. As the global long-term growth rate *M*_*i*_ − *V*_*i*_*/*2 equals zero, the difference *M*_*i*_ − *V*_*i*_ equals the negative quantity −*V*_*i*_*/*2. Hence, in patches acting as buffers, the local mean growth rate is negative, and, consequently, these patches are long-term, deterministic sinks i.e. 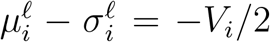 and 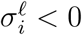 implies that 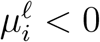.

#### Whenever multiple patches are occupied, all are long-term sinks

Provided there is some spatial asynchrony in the environmental fluctuations experienced by species *i* (i.e. Σ_*i*_ is non-degenerate), the patch-selection strategy of species *i* at the coESS exhibits a fundamental dichotomy (Proposition D.1 in Appendix D): either species *i* only occupies one patch, or it occupies multiple patches and the local long-term growth rates are negative in these occupied patches. Thus, if species *i* occupies multiple patches at the coESS, then all of its populations are long-term sink populations despite it persisting globally. This occurs as species playing the coESS exhibits a form of spatial bet-hedging that results in the global stochastic growth rate being greater, specifically zero, than the local stochastic growth rates (see discussion).

Evolution for occupying a single patch in the landscape only occurs when the other patches are long-term sinks. Moreover, these sinks must lead to sufficiently negative long-term growth rates or exhibit similar environmental fluctuations as the occupied patch. More precisely, our coESS condition (9) (Proposition D.2 in Appendix D) implies that only patch *ℓ* is occupied if

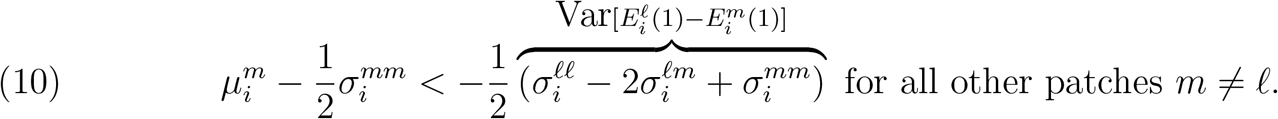

To better understand (10), consider the case where the environmental fluctuations for species *i* in all patches have variance *σ*^2^ and spatial correlation *ρ*, i.e.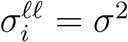 for all *ℓ* and 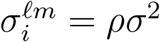 for *ℓ* ≠ *m*. Then, condition (10) requires that the long-term growth rates in the unoccupied patches *m* are less than −*σ*^2^(1 − *ρ*). Hence, selection for only occupying patch *ℓ* is greatest when the environmental fluctuations across patches are strongly, positively correlated (*ρ* ≈ 1).

### Applications to antagonistic interactions

Antagonistic interactions, such as the interactions between predators and their prey or between competing species, can result in reciprocal selection pressures. Here, we investigate how this reciprocal evolution can either lessen the antagonism by selecting for divergent choices in patch use, or enhance the antagonism by selecting for convergent choices in patch use.

### Predator-prey coevolution

#### A general model

We begin with a predator-prey system where *x*_1_ and *x*_2_ are the global densities of the prey and predator, respectively. The intrinsic per-capita growth rate of the prey in patch *ℓ* is *r*^*ℓ*^. The predator is a specialist on the prey with attack rates, conversion efficiencies, and per-capita death rates in all patches equal *a, c*, and *d*, respectively. As the predator is a specialist, it can not persist in the absence of the prey species. To ensure stability of the predator-prey dynamics, we assume that both species experience weak intraspecific competition with strength *α* ≈ 0. Under these assumptions, the Lotka-Volterra model takes on the form

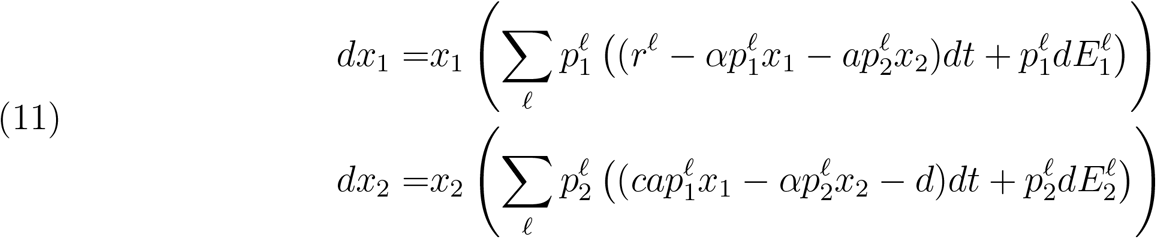

In Appendix E, we derive criteria which characterize when the species coexist globally, and find explicit expressions for the mean densities of the species at stationarity and the invasion growth rates 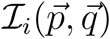. Here, we focus on two cases: a two-patch sink-source system and an environmental gradient for the prey.

#### A two-patch source-sink system

Consider a landscape where the prey has a source habitat (patch 1 with *r*^1^ = *r*_source_ *>* 0) and a sink habitat (patch 2 with *r*^2^ = − *r*_sink_). The sink habitat is low quality but exhibits minimal environmental fluctuations 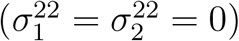, while the predator and prey experience environmental fluctuations in the source habitat with variances *v*_prey_ and *v*_pred_, respectively.

First, we study the effects of environmental stochasticity on the species individually and then collectively. If only the prey experiences environmental stochasticity in the source patch (*v*_prey_ *>* 0 and *v*_pred_ = 0), then the coESS has neither species occupying the sink patch for low values of *v*_prey_, the prey occupying both patches at intermediate values of *v*_prey_, both species occupying both patches at higher values of *v*_prey_, the predator no longer occupying the source patch at even higher values of *v*_prey_ and, finally, extinction of both species when *v*_prey_ is too high (Figure 1A,B).

**Figure 1.**
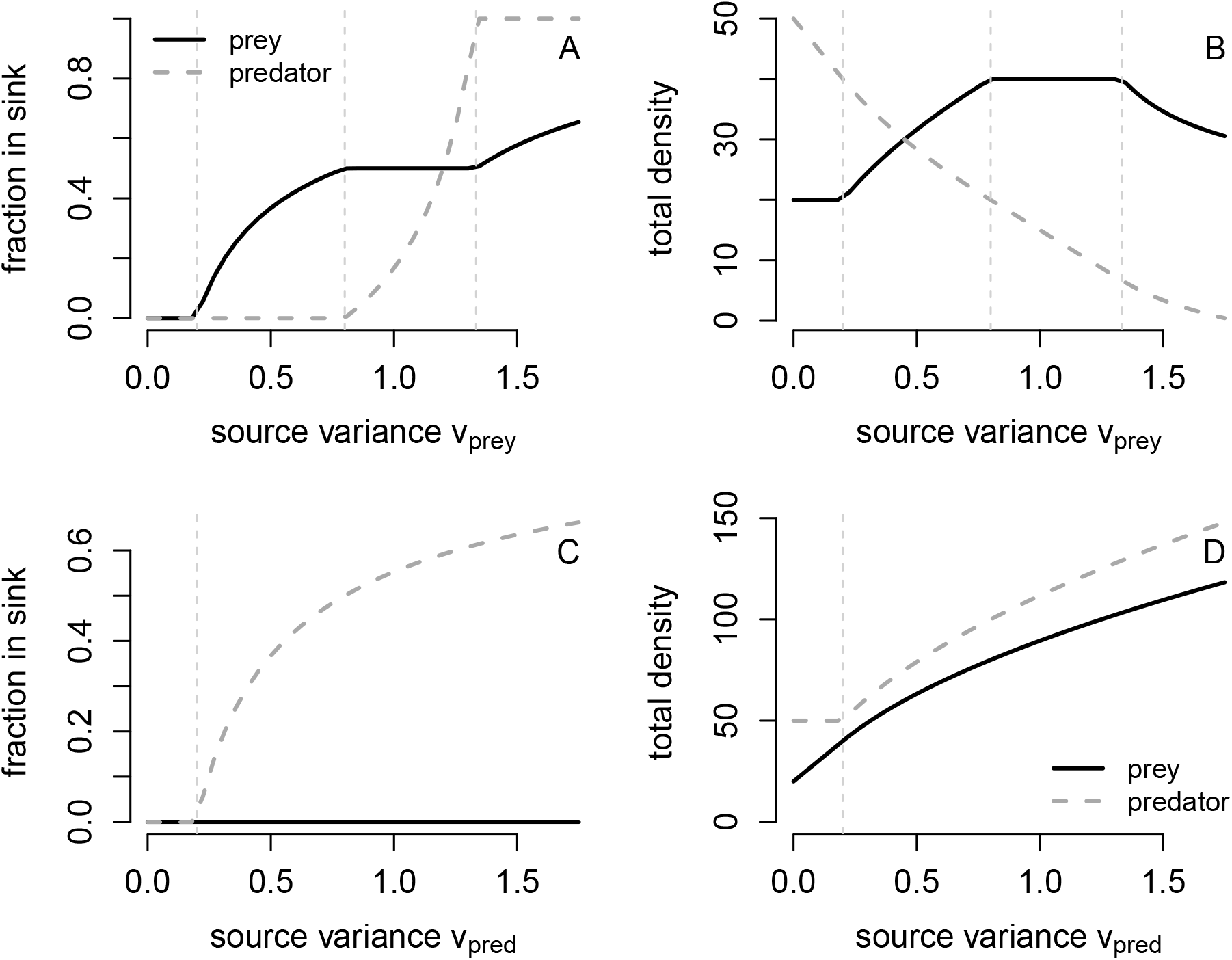
The coESS for patch choice and the mean global densities for a predator-prey system with a source patch and a sink patch. In (A) and (B), the prey experience environmental fluctuations in the source patch with variance *v*_prey_. In (C) and (D), the predator experiences environmental fluctuations in the source patch with variance *v*_pred_. Solid and dashed thick lines correspond to numerically estimated solutions by simulating (8) for 2, 000 time steps. Dashed thin, vertical lines correspond to the analytic conditions for changes in patch use presented in text. Parameter values: *r*_source_ = 0.5, *r*_sink_ = 0.1, *d* = 0.1, *a* = 0.01, *c* = 0.5, intraspecific competition coefficient of 0.000001 for both species.

More specifically, when the environmental variance in the source patch is sufficiently low relative to the rate of loss in the sink patch (*v*_prey_ *<* 2*r*_sink_), inequality (10) implies the prey only occupies the source patch. As the coESS condition (9) requires that the growth rate of the predator is equal to zero in any patch it occupies, the predator also only occupies the source patch. When the environmental variance in the source patch exceeds 2*r*_sink_ but lies below 8*r*_sink_, the coESS conditions (9) imply that the fraction of prey living in the sink patch equals

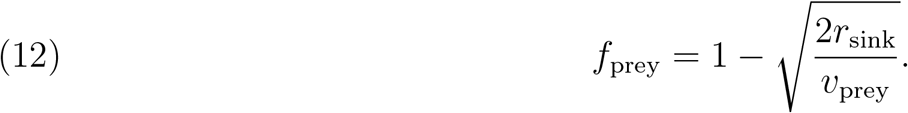

Equation (12) implies that the fraction of prey in the sink patch increases with the environmental variance *v*_prey_ in the source patch (Figure 1A). As the predator regulates the mean prey density in the source patch to the predator’s break-even point 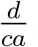, selection for prey using the sink patch results in higher mean prey densities (Figure 1B, Appendix E). These trends continue with increasing variation in the environmental fluctuations until 50% of the prey make use of the sink patch.

When the environmental stochasticity in the source patch selects for the prey being equally distributed between the patches (i.e 8*r*_sink_ ≤ *v*_prey_ ≤ 8*r*_source_*/*3 and 3*r*_sink_ ≤ *r*_source_, Appendix E), the predator evolves to use both patches. The fraction of predator using the sink patch equals

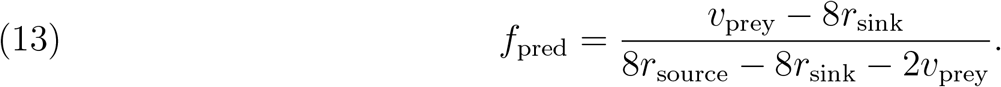

Equation (13) implies the fraction *f*_pred_ of predators in the sink patch increases with the environmental stochasticity *v*_prey_ experienced by the prey. As the predator does not directly experience environmental stochasticity, it regulates the prey density, on average, to its break-even point in both patches. Hence, for this range of environmental variation, the mean global density of the prey is twice as high as when both species only reside in the source patch (Figure 1B). In contrast, the predator’s mean global density decreases with increasing variation in the environmental fluctuations (Figure 1B).

When the variation in the environmental fluctuations are sufficiently large 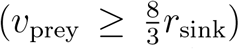 but not so large as to cause extinction, the predator evolves to only use the sink patch (i.e. *f*_pred_ = 1) while the prey continues to use both patches. When this occurs, the fraction of prey making use of the sink patch equals (Appendix E)

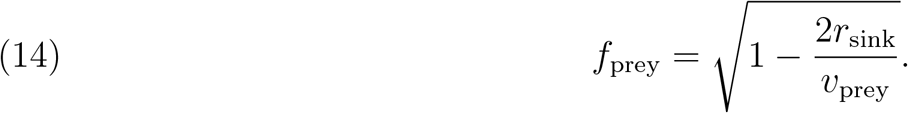

Equation (14) implies the fraction of prey in the sink patch continues to increase with increasing variation of the environmental fluctuations. In contrast, the mean global prey density decreases with the environmental variance (Figure 1B, Appendix E). When the environmental fluctuations are sufficiently strong, 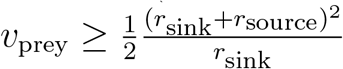, both species go extinct (not shown in Figure 1).

When only the predator experiences environmental stochasticity in the source patch (*v*_pred_ *>* 0 and *v*_prey_ = 0), the coESS condition (9) implies that the prey’s growth rate is zero in all occupied patches. Consequently, prey playing the coESS only occupy the source patch. Despite a victimless sink patch, the predator evolves to occupy the sink patch whenever the environmental variance in the source patch is sufficiently great i.e. *v*_pred_ *>* 2*d*. Under these circumstances, the fraction of predators using the sink patch equals

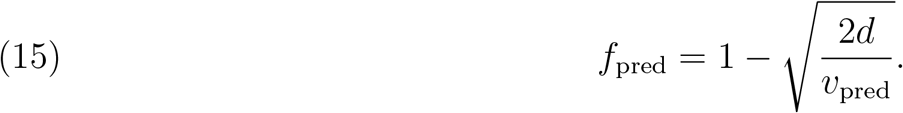

Equation (15) implies greater environmental variation in the source patch selects for greater use of the sink patch (Figure 1C). When the predator makes use of the sink patch, the mean global density of both species playing the coESS increases with the environmental variance in the source patch (Figure 1D, Appendix E). Intuitively, as the environmental fluctuations increase, the predator has a higher break-even prey density in the source patch that determines the mean prey density. These higher prey densities, in turn, lead to higher mean predator densities.

Figure 2 illustrates the effects of simultaneous environmental variation on both species on the coESS. Consistent with the predictions from varying environmental stochasticity for only one species, high environmental variation for the predator selects for victimless sinks when variation for the prey is sufficiently low, and selects for both species using both patches when this variation is sufficiently high. In contrast, low environmental variation for the predator selects for enemy-free sinks, both species using the sink patch, and enemy-free sources with increasing levels of environmental variation for the prey.

**Figure 2.**
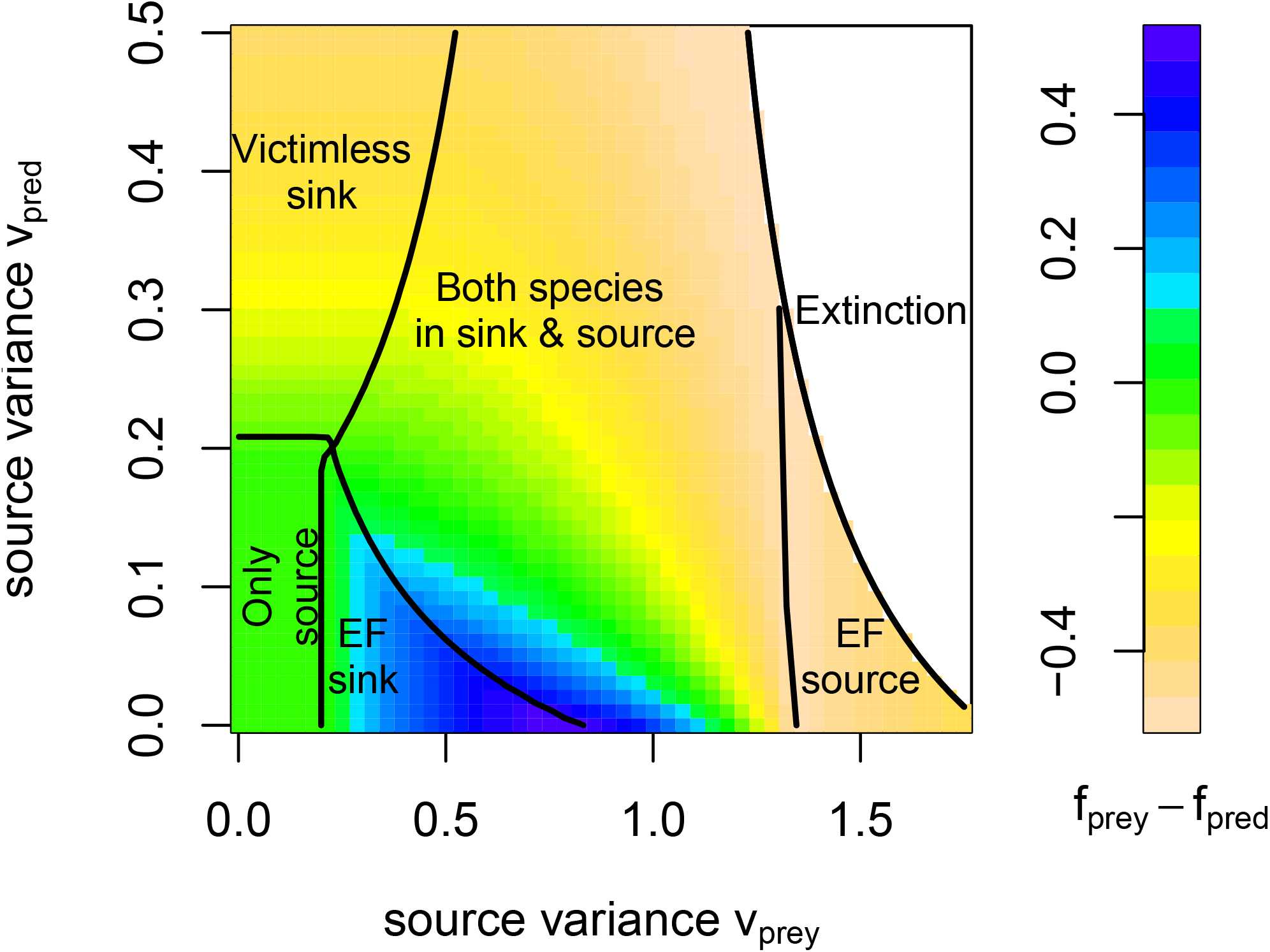
How simultaneous environmental fluctuations for predator and prey selects for enemy-free (EF) patches, victimless sinks, and predator-prey interactions in sink patches. White region corresponds to environmental variation that leads to extinction of both species. Colors represent the difference in the fraction of prey and predators in the sink patch. Parameters as in Figure 1.

#### Patch-selection along an environmental gradient

Using our numerical algorithm, we examined patch-selection along a gradient of environmental fluctuations. Along this gradient, the environmental variance experienced by both species varies in a Gaussian manner (Figure 3A). In the center of the landscape where environmental fluctuations are strongest, the patches are long-term, stochastic sinks (patches between dashed vertical lines in Figure 3). When the environmental fluctuations are spatially uncorrelated, all patches are occupied by both species but the species exhibit contrary choices: the two species exhibiting negatively-correlated patch selection strategies. (Figure 3B). Spatial correlations in the environmental fluctuations selects for both species exhibiting reduced preferences and lower densities in the most central patches (Figure 3C,D). When these spatial correlations are sufficiently strong, neither species occupies the central patches consistent with the general prediction of approaching an ideal-free distribution (Figure 3D). For intermediate spatial correlations, only the predator avoids the central patches creating enemy-free sinks (Figure 3C).

### Exorcising the ghost of competition past

#### A general model

To understand how spatial-temporal heterogeneity selects for spatial distributions of competing species, we examine a model of two species competing in *k* patches. This analysis provides insights into when competitors evolve to never occupy the same patches i.e. the ghost of competition past (Lawlor and Maynard Smith, 1976; Connell, 1980), or evolve to co-occur within patches i.e. exorcising the ghost. For competitor *i* in the model, its intrinsic rate of growth in patch *ℓ* is 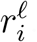. The average per-capita growth rates decrease linearly with the local density of both competitors. Namely, the average per-capita growth rate of species *i* in patch *ℓ* equals 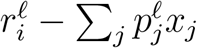. Thus, the global dynamics are

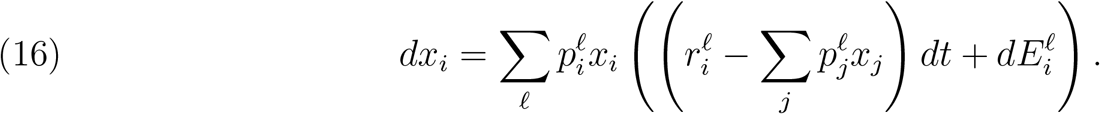

These competitors, in general, can not coexist locally: the species with the larger intrinsic stochastic growth rate 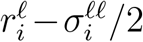 in patch *ℓ* excludes the other species in the absence of immigration. Global coexistence, however, is possible in a multi-patch landscape. In Appendix F, we derive criteria which characterize when the species coexist globally, the mean equilibrium densities of the species at the associated stationary distribution, and explicit expressions for the invasion rates of mutant strategies against resident strategies. Here, we focus on two cases: two species in a spatially symmetric landscape and three species living along an environmental gradient.

#### Competition along a symmetric landscape

Imagine a symmetric landscape with an even number of patches where each species has the competitive advantage in half of the landscape. The average per-capita growth rate for the species *i* in the patches where it is competitively superior equals 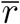 and equals 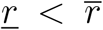 in the patches where it is competitively inferior. Each species experiences spatially uncorrelated, environmental fluctuations with local variance *v*. Our analysis reveals that there is critical level of environmental variance, below which the species never occupy the same patch, and above which this ghost of competition past is exorcised. The critical level of variance depends both on the number of patches in the landscape and the fitness difference of the competitors.

To see why these conclusions hold, the symmetry of the landscape implies that any coESS for the competing species satisfies that a fraction *f* of individuals use the patches where they have competitive disadvantage (sink patches), and the complementary fraction 1 − *f* use the patches where they have a competitive advantage (source patches). At the coESS (see Appendix F for details), each species only uses the source patches (i.e. *f* = 0) if

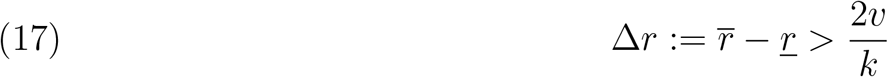

Equation (17) implies that if the environmental variance *v* is too low relative to the number of patches, or the fitness difference Δ*r* is too large, then evolution selects for the competitors to be spatially segregated i.e. the ghost of competition past prevails (left-hand and right-hand sides of Fig. 4A,C, respectively).

**Figure 3.**
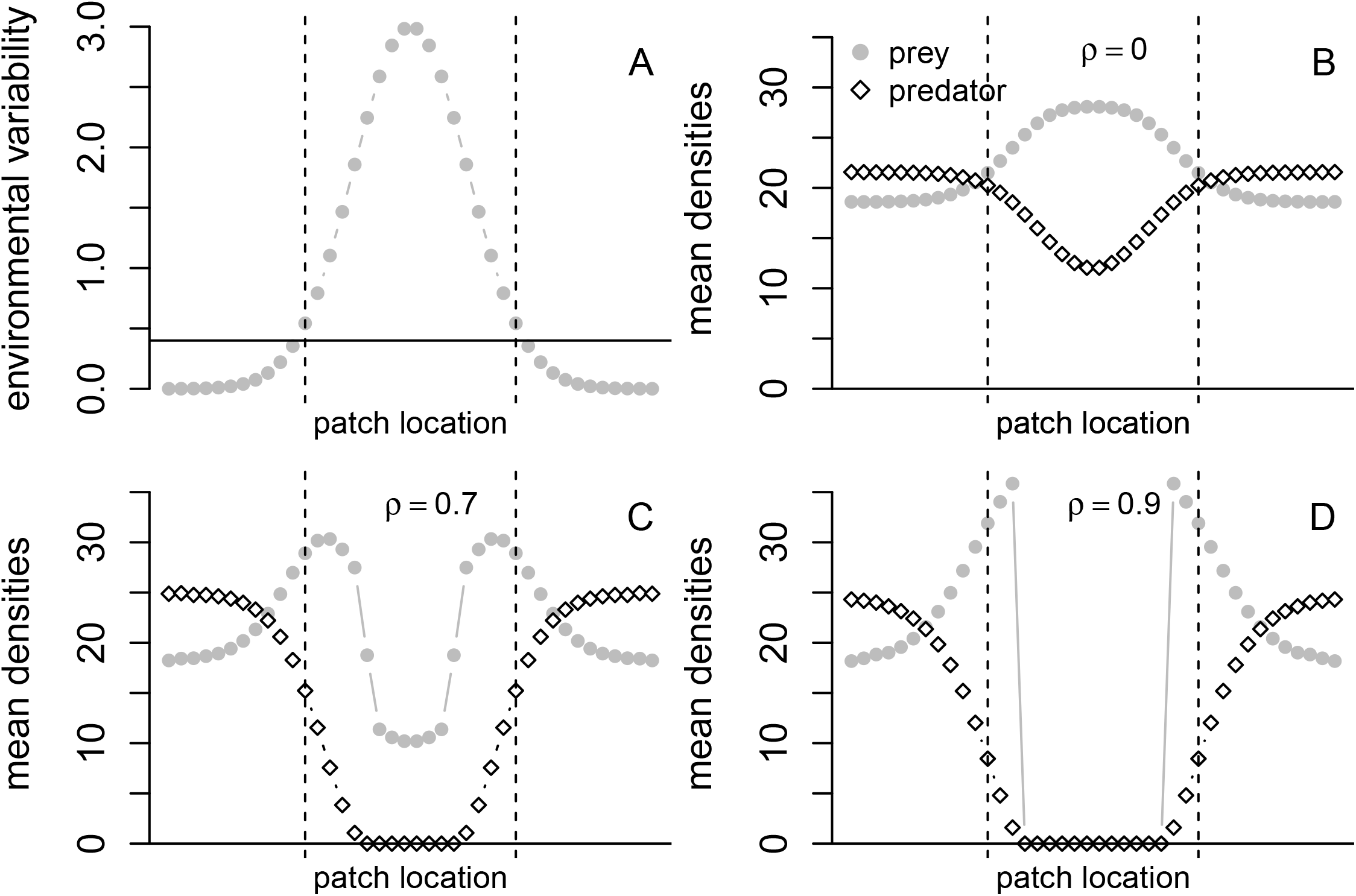
CoESS for predator-prey interactions along a spatial gradient of environmental fluctuations. Both species experience environmental fluctuations whose strength decay from a central location in the landscape (panel A). Horizontal line in A corresponds to twice the prey’s intrinsic rate of growth (2*r*^*ℓ*^). Patches between the vertical dashed lines are long-term sources for the prey. Patches outside of the vertical dashed lines are long-term, stochastic sinks for the prey. The spatial correlation *ρ*^| *ℓ*−*m*|^ between two patches *ℓ* and *m* decays with distance where *ρ* is the spatial correlation between two neighboring patches i.e. |*ℓ* −*m*| = 1. The mean densities of the predator (white diamonds) and prey (gray dots) are plotted for the coESS at three levels of correlation *ρ* (panels B-D). Parameters: *n* = 2 species, *k* = 40 patches, *r*^*ℓ*^ = 0.1, *a*^*ℓ*^ = 0.01, *c*^*ℓ*^ = 0.5, *d*^*ℓ*^ = 0.1 for all *ℓ*, 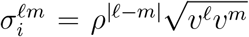 where *v*^*ℓ*^ = 3 exp(−(*z*^*ℓ*^)^2^) for *z* = 6*/ℓ* − 3 and 1 ≤ *ℓ* ≤ 40, *ε* = 0.00001.

**Figure 4.**
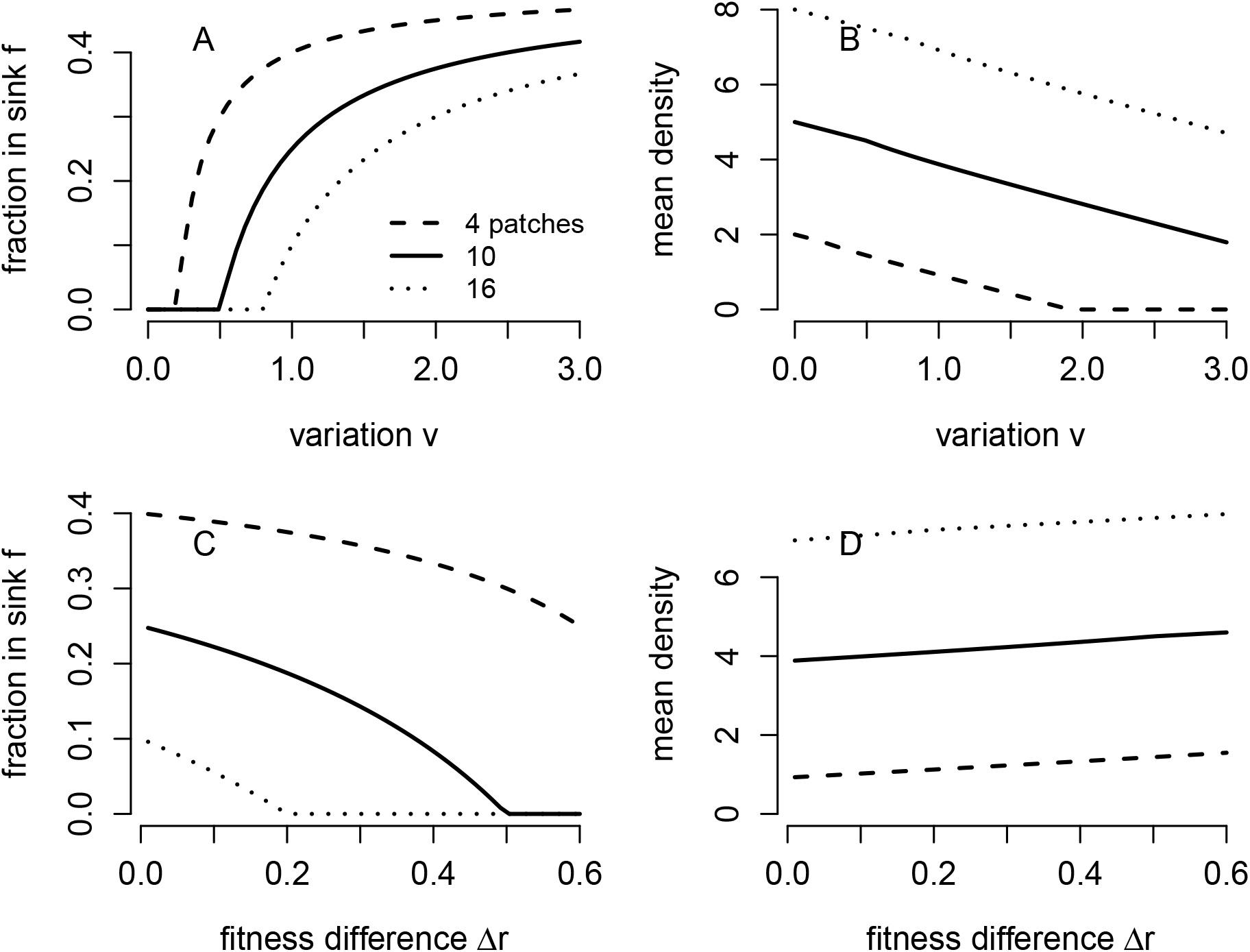
The effects of environmental variation *v*, fitness differences 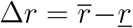, and number *k* of patches on the fraction of competitors using the patches in which they are the inferior competitor (A,C) and the mean global density of each competitor (B,D). Parameters: 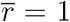 in all panels, *r* = 0.9 in (A,B), *r* = 1 − Δ*r* and *v* = 1 in (C,D).

When environmental fluctuations are sufficiently large (i.e. inequality (17) is reversed), evolution selects for both species to use both patch types with

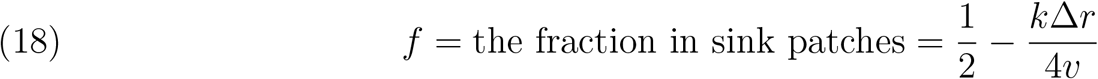

i.e. the ghost of competition past is partially exorcised. Equation (18) implies that the majority of individuals (i.e. 1 − *f >* 1*/*2) occupy their source patches. However, for smaller fitness differences Δ*r* or higher levels of environmental variation *v*, evolution selects for both species to be spread more equally across the patches (i.e. right-hand and left-hand sides of Fig. 4A,C, respectively).

Whether sink patches are occupied or not, the mean global density of each competitor equals

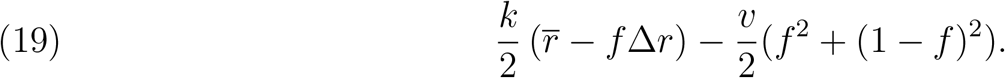

When the competitors are spatially segregated (i.e. *f* = 0), the equilibrium density 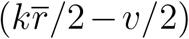 is a decreasing linear function of the environmental variation (Fig. 4B). Selection for the use of both patch types (i.e. *f >* 0) results in a nonlinear, accelerated, negative response of mean density to increasing environmental variation.

#### Three species competition along an environmental gradient

Using our numerical algorithm, we also explored coevolution of patch selection for three competing species along an environmental gradient (Figure 5). Along this gradient, the species differed in their intrinsic rates of growth and experienced the same amount of environmental stochasticity (Figure 5A). In the absence of environmental fluctuations, the coESS corresponds to an ideal-free distribution resulting in each species only occupying the patches in which their intrinsic stochastic growth rates are positive and in which they are competitively superior (Figure 5B). Notably, the species are spatially segregated and occupy no sink patches. In the presence of environmental fluctuations, each patch is occupied by at least two species – the ghost of competition past is partially exorcised (Figure 5C). Moreover at the center and edges of the landscape, all three species co-occur. At the edges, the co-occurring species are long-term, determinsitic sink populations.

**Figure 5.**
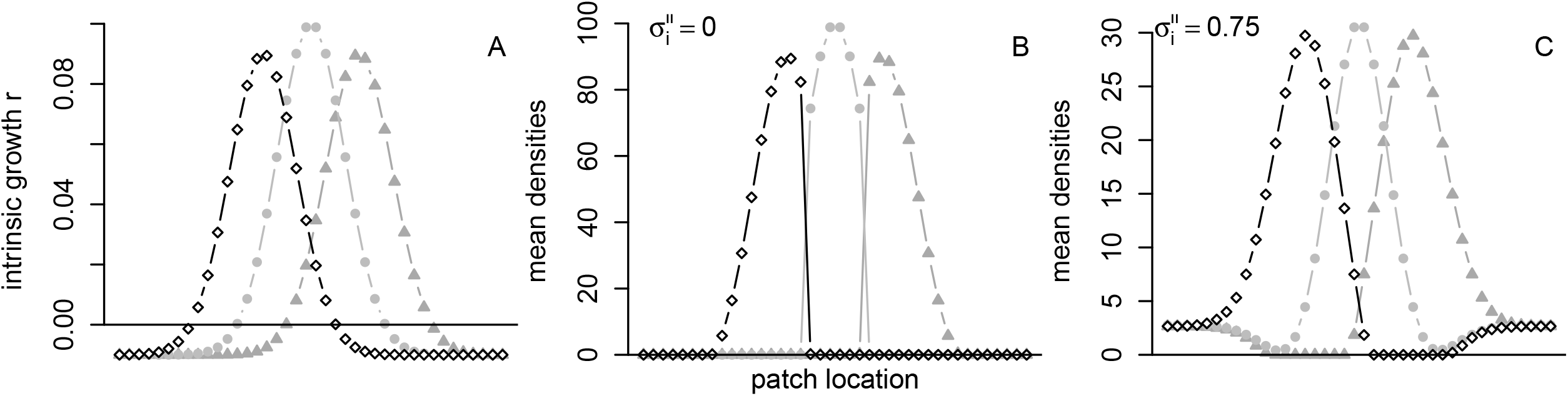
CoESS for three competing species along an environmental gradient. In A, the environmental variation in the average intrinsic rates of growth 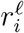 for all three species. In B and C, the mean densities of all three species playing the coESS in the absence (B) and presence (C) of environmental stochasticity. Parameters: *n* = 3 species, *k* = 40 patches, 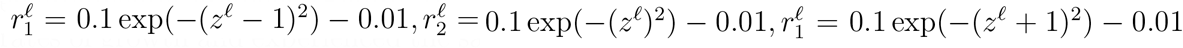 for *z*^*ℓ*^ = 8*/ℓ* − 4 and 1 ≤ *ℓ* ≤ 40, 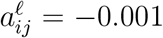 for all *i, j, ℓ*, and Σ_*i*_ = *σ*^2^Id with *σ*^2^ = 0 in B and = 0.75 in C.

## Discussion

Unlike classical ideal-free theory that predicts selection against sink populations under equilibrium conditions (Fretwell and Lucas, 1969; Křivan et al., 2008), we find that coevolution in fluctuating environments often selects for metacommunities consisting entirely of long-term sink populations. The difference stems from what selection equalizes across the occupied landscape under equilibrium versus non-equilibrium conditions. Under equilibrium conditions, the percapita growth rates of all populations are equal to zero in occupied patches and negative in unoccupied patches. Hence, there are no sink populations. In landscapes with environmental stochasticity, coevolution of patch-selection no longer equalizes the mean per-capita growth rates in occupied patches. Instead, the differences between the means of the local per-capita growth rates and the covariance between the local and global fluctuations are equalized. Whenever there is evolution for occupying multiple patches, equalizing these differences sacrifices higher mean growth rates for lower variances of these growth rates and results in negative long-term growth rates in all occupied patches. This spatial bet-hedging can be understood through the lens of the Modern Portfolio Theory (MPT) of economics (Markowitz, 1952, 1991; Rubinstein, 2002; Markowitz, 2010), as we discuss below. The resulting sink populations may be conditional due to the presence of antagonistic interactions, or unconditional. In particular, environmental stochasticity coupled with predator-prey interactions can select for enemy-free sinks and victimless sinks (Jeffries and Lawton, 1984). Alternatively, environmental stochasticity coupled with competitive interactions can select for conditional sink population by partially exorcising the ghost of competition past (Lawlor and Maynard Smith, 1976; Connell, 1980; Schmidt et al., 2000).

### Relation to Modern Portfolio Theory

Modern Portfolio Theory (MPT) (Markowitz, 1952, 1991, 2010) is a Nobel prize winning framework for assembling a portfolio of financial investments. In this framework, each investment has a mean return and some level of risk characterized by the variance in the return. In our context, investments correspond to patches, mean returns of investments correspond to mean local growth rates 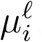, and risks correspond the local environmental variances 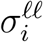(Figure 6A). A patch-selection strategy (Figure 6B) 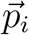 corresponds to a portfolio of investments i.e. a fixed allocation of assets across the investments. The global mean growth rate *M*_*i*_ and the global environmental variance *V*_*i*_ of a patch-selection strategy corresponds to the mean return and variance of a portfolio of investments, respectively. MPT assumes that investors prefer portfolios with high mean returns and avoid those with high risk. As investments with high mean returns often are high risk investments, investments often exhibit a mean-variance trade-off (Zivot, 2017). In our context, however, patches, themselves, need not exhibit a mean-variance trade-off; highquality patches may consistently support high mean growth rates and low-quality patches may be highly variable. Instead, the mean-variance trade-off stems from the long-term growth rate increasing with the mean but decreasing with the variance (*M*_*i*_ − *V*_*i*_*/*2 or 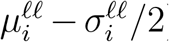), the corner stone of bet-hedging theory in evolutionary biology (Cohen, 1966; Stearns, 2000; Childs et al., 2010). Markowitz (1952) showed that for a given level of risk *V*, there are portfolios maximizing the mean return *M* for that level of risk. These optimal portfolios can be solved for only using matrix algebra (Zivot, 2017). The curve of means and variances determined by these portfolios is the efficient frontier (solid curves in Figure 6C,D). Markowitz (1952) showed that the efficient frontier also corresponds to the portfolios minimizing an investor’s risk for a given mean return. In our context, the coESS patch-selection strategy must be a point on the efficient frontier (white circles in Figure 6C,D). Indeed, if it wasn’t, there would be an alternative patch-selection strategy either providing lower risk for the same mean return or providing a greater mean return for the same amount of risk. In either case, such a patch-selection strategy would have a higher global long-term growth rate *M*_*i*_ − *V*_*i*_*/*2 than residents playing the coESS – contradicting the definition of a coESS. By lying on the efficient frontier, the coESS typically sacrifices a higher global mean growth rate for a lower global variance (all the local mean growth rates are greater than the mean growth rate of the coESS in Figure 6C,D). Hence, the coESS typically is a bet-hedging strategy (Childs et al., 2010). The only exception occurs when the mean growth rates are equal in all occupied patches. Only then, can the coESS can reduce risk without reducing the mean. Furthermore, as the global long-term growth rate *M* − *V/*2 is zero, the coESS corresponds to the point on the efficient frontier that is tangent to the *M* = *V/*2 line (see white circle and dashed lines in Figures 6C,D) – in economic terms, a tangent portfolio associated with a risk-free, zero return asset (Zivot, 2017). Whenever multiple patches are occupied at the coESS, all the points in the mean-variance plane corresponding to the individual patches (i.e. individual investments) lie below efficient frontier and, consequently, have negative local long-term growth rates (see shaded circles in Figures 6C,D). Hence, all patches are long-term sinks.

**Figure 6.**
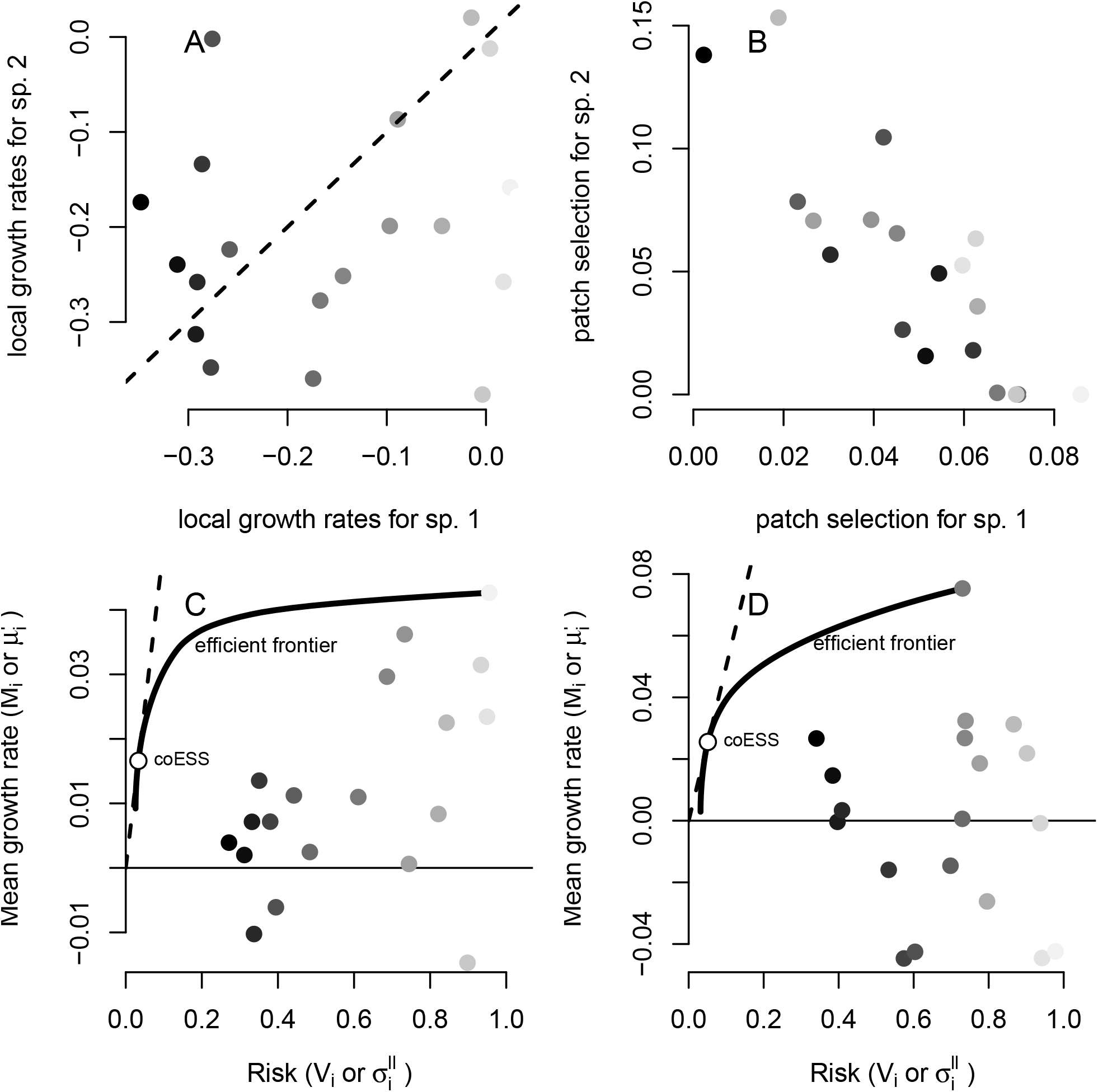
The coESS of patch-selection through the lens of Modern Portfolio Theory (MPT). The comparison considers two competing species in a 20 patch landscape with differing local long-term growth rates (panel A). These local long-term growth rates determine local competitive superiority. The coESS for patch-selection (panel B) exhibits a negative correlation between the competitors. At the coESS, the local mean growth rates and local variances are plotted in the mean-variance plane for each of the species (panels C, D). The efficient frontier (see text) is plotted as a solid line, the global means and variances associated the coESS are plotted as a white circle, and the dashed line is where the global stochastic growth rate equals zero.

### Ghosts of past and present antagonisms

Patches free of antagonistic interactions may reflect habitat selection in response to past antagonisms within a patch or present antagonisms in other patches. Spatial heterogeneity can select for ghosts of competition past or enemy-free space by creating spatial mosaics of environmental conditions favoring one species over another (Lawlor and Maynard Smith, 1976; Rosenzweig, 1981, 1987; Schreiber et al., 2000; Schreiber and Vejdani, 2006). Our results demonstrate that environmental stochasticity can reverse or amplify these outcomes.

We find that environmental stochasticity can partially exorcize the ghost of competition past. Under equilibrium conditions, ideal-free competitors only occupy habitat patches in which they are competitively superior (Lawlor and Maynard Smith, 1976; Connell, 1980; Rosenzweig, 1981, 1987). When environmental fluctuations are sufficiently large relative to fitness differences between the competing species, we find that there is selection for competitors occupying patches in which they are competitively inferior – conditional sink populations. Using computer simulations, Schmidt et al. (2000) observed a similar exorcism of the ghost of competition past for discrete-time, two patch models of two competing species. Our results provide an analytical extension of their work to any number of patches and any number of competing species. Moreover, we find that along environmental gradients, environmental stochasticity can select for several competitors occupying sink habitat patches at the edges of these gradients i.e. unconditional sink populations. This simultaneous selection for conditional and unconditional sink populations can result in complex species distributions along an environmental gradient e.g. species disappearing and reappearing along the gradient. Similar complex patterns have been observed along elevational gradients in birds (Noon, 1981; Campos-Cerqueira et al., 2017). For example, on Camel Hump’s Mountain, the hermit thrush (*Catharus guttatus*) is most common at lower (400m) and higher (800m) elevations but is less common at intermediate (600m) elevations (Noon, 1981). Whether or not environmental fluctuations play a role in these empirical patterns remains to be understood.

Our results highlight how environmental fluctuations can select for enemy-free space in two ways. First, environmental fluctuations in higher quality habitat may select for prey using lower quality habitats and, thereby, create unconditional sink populations. If these lower quality habitats only support low prey densities and the predators are less sensitive to the environmental fluctuations, then the predators may evolve to only occupy the higher quality habitat. Thus, the lower-quality habitat becomes enemy-free space. In this case, the spatial bet-hedging by the prey creates the enemy-free space. An alternative pathway to enemy-free space occurs when both species are sensitive to the environmental fluctuations and the intensity of these fluctuations vary across the landscape. This spatial variation can select for contrary choices – prey selecting patches with greater risk and predators selecting patches with lower risk. This form of selection for enemy-free space is a stochastic analog of fixed spatial heterogeneity selecting for contrary choices (Fox and Eisenbach, 1992; Schreiber et al., 2000, 2002; Schreiber and Vejdani, 2006): prey select lower quality patches to lower the reproductive success of their predators and predators selecting high quality patches to maximize their per-capita reproductive success.

### Eco-evolutionary hydra effects

Our results on predator-prey coevolution illustrate how environmental stochasticity drive hydra effects over evolutionary time. Named after the mythological beast who grew two heads to replace a lost head, a hydra effect occurs when a population’s mean density increases in response to an increase in its per-capita mortality rate (Abrams, 2009; Sieber and Hilker, 2012; Cortez and Abrams, 2016). For example, Abrams (2009) found that increasing the per-capita mortality rate of a predator with a type II functional response could lead to long-term increase the predator’s mean density. Similar to increasing per-capita mortality rates, increasing environmental stochasticity (*σ*^2^) reduces stochastic per-capita growth rates (from *µ* to *µ* − *σ*^2^*/*2) and, consequently, often has negative impacts on average population densities in the long-term (Lande et al., 2003). For example, for Lotka-Volterra predator-prey dynamics with environmental stochasticity, increasing environmental stochasticity experienced by the predator decreases its average density in the long-term (May, 1975) (see, also, the mean density 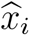 expressions in Appendix E). However, we find that if there is sufficient time for the patch-selection strategies of the predator and prey to evolve to their new coESS, then increasing environmental stochasticity may cause the average predator density to increase (Figure 1D). Intuitively, sufficiently high environmental stochasticity can select for a predator to hedge its bets by occupying an environmentally stable, but low quality patch. This spatial bet-hedging lowers the predation pressure on the prey in the source habitat resulting in an increase of its average density and a corresponding increase in the average predator density. We found similar hydra effects when the prey experiences increasing levels of environmental stochasticity (Figure 1B).

### Caveats and future directions

To simultaneously confront the complexities of species coevolution in a spatially and temporally-variable environment, we made several simplifying assumptions. Relaxing these assumptions provide significant challenges for future research. Most importantly, our framework assumes that species do not assess or respond to temporal changes in habitat quality; they exhibit a fixed spatial distribution. While this assumption is a good first-order approximation frequently made in the theoretical literature (Hassell et al., 1991; van Baalen and Sabelis, 1993; Holt, 1997; Jansen and Yoshimura, 1998; Schmidt et al., 2000; Schreiber et al., 2000; Schreiber and Vejdani, 2006; Schreiber, 2012), spatial distributions typically vary in time in response to environmental fluctuations. Hence, a major challenge for future work is developing methods to study the evolution of dispersal rates in spatially explicit and temporally variable landscapes. While there has been extensive analytical and numerical work on this question for single species (Levin et al., 1984; McPeek and Holt, 1992; Hutson et al., 2001; Ronce, 2007; Cantrell and Cosner, 2018; Cantrell et al., 2021), much less work exists for interacting species (see, however, Lion et al. (2006); Schreiber and Saltzman (2009); Lion and Gandon (2015)).

As most ecological communities reside in spatially and temporally variable environments, we might expect that many of our qualitative predictions occur in nature. However, as noted by Urban et al. (2020), “to date, few empirical studies completely evaluate eco-evolutionary interactions in space through field manipulations, and even fewer test for underlying mechanisms.” Ideally, to test the theory presented here, one would collect data on spatial and temporal variation in species demographic rates, interaction strengths, and densities over sufficiently long time frames. Alternatively, one could evaluate if key ingredients of conditions and consequences hold for interacting species occur over some environmental gradient or patchy landscape. For example, our results predict that greater environmental fluctuations will select for greater range overlap of competing species. Hence, one could attempt a meta-analysis similar to Urban et al. (2020) to evaluate across multiple competitive metacommunities whether such a correlation exists, ideally controlling for the degree of spatial heterogeneity across the landscape. Such analyses could identify whether or not reciprocal selection among interacting species give rise to some of outcomes predicted by our theory.

### In conclusion

In conclusion, our work provides one theoretical approach to studying how spatial heterogeneity, temporal variation, and species interactions drive the evolution of habitat choice. In this framework, stochastic sink populations are the norm, not the exception due to species using space to hedge their bets. For antagonistic species interactions, this bet-hedging can select for predators seeking refuge in prey-free space and can exorcise the ghost of competition past. We anticipate that applying these methods to models with more species will reveal new surprises.

## Appendix A. Stochastic Coexistence

In the past decade, there has been a substantial development of new methods for determining coexistence and extinction for stochastic difference equations and stochastic differential equations (Schreiber et al., 2011; Benaïm and Lobry, 2016; Hening and Nguyen, 2018*a*; Benaïm, 2018; Benaïm and Schreiber, 2019; Hening et al., 2021). All of these methods all rely on invasion growth rates: the long-term average per-capita growth rate of a species when rare. Importantly, Hening and Nguyen (2018*a*); Hening et al. (2021) developed methods that apply to stochastic Lotka-Volterra systems of the form

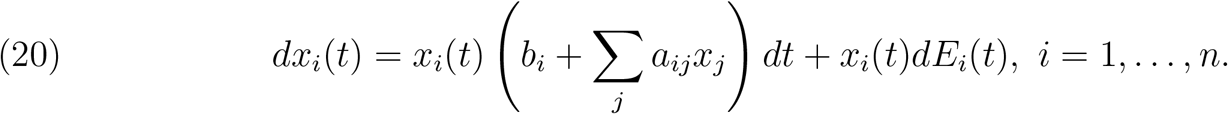

Here **E**(*t*) = (*E*_1_(*t*), …, *E*_*n*_(*t*))^*T*^ = Γ^⊤^ **B**(*t*), Γ is a *n × n* matrix such that Γ^⊤^Γ = ∑ = (*σ*_*ij*_)_*n*×*n*_ and **B**(*t*) = (*B*_1_(*t*), …, *B*_*n*_(*t*)) is a vector of independent standard Brownian motions. Our main model (3) can be written in the form of (20) by matching the drift coefficients and the covariances - see for more details Section 3 in Evans et al. (2015). Specifically, (3) is equivalent to (20) by making the following definitions

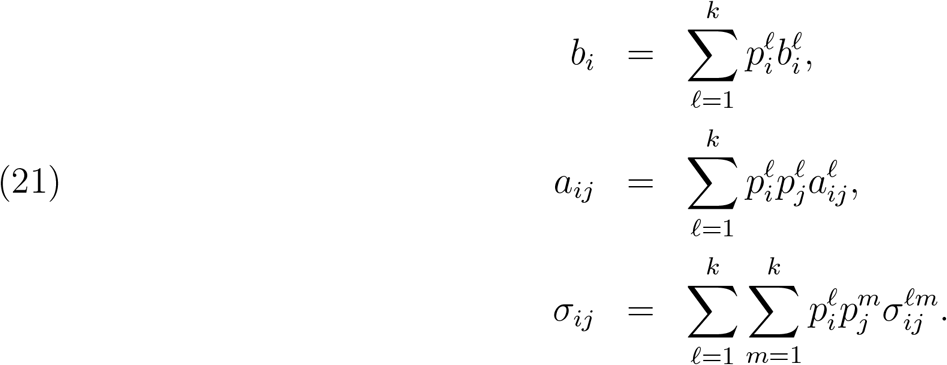

To introduce the invasion growth rates, suppose we introduce species *i* at an infinitesimally small density when the dynamics of the remaining species follows the stationary distribution *μ*(*d***x**). Applying Itô’s lemma to log *x*_*i*_ in (20), we get the average per-capita growth rate of the invading species *x*_*i*_ at time *t* is

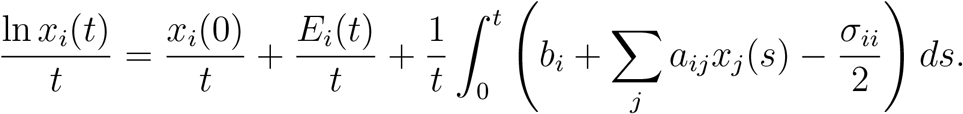

In the long run, the term 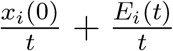 is negligible and the integral term can be approximated by averaging over the stationary distribution:

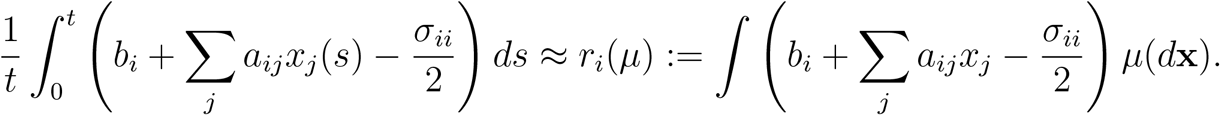

As a result, it is natural to define the long-term growth rate of species *i* into the community of species following the stationary distribution *μ*(*d***x**) as

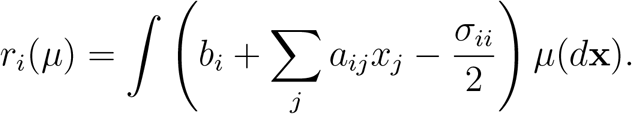

When *r*_*i*_(*μ*) *<* 0, species *i* tends to decline at an exponential rate. When *r*_*i*_(*μ*) *>* 0, species *I* tends to increase at an exponential rate.

Following Chesson (1982) we say species *i* is stochastically persistent if with high probability the species is bounded away from low densities. We say there is stochastic persistence if all species are persistent. According to whether *r*_*i*_(*μ*) *>* 0 or *r*_*i*_(*μ*) *<* 0, if 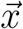 spends a long time close to the support of *μ*, its *i*th component will get repelled from or attracted towards the face *x*_*i*_ = 0. Under some natural assumptions, it was shown in Hening and Nguyen (2018*a*) that if every stationary distribution *μ*(*d***x**) on the extinction set is a repeller, i.e. max_*i*_ *r*_*i*_(*μ*) *>* 0 then the species persist and, under certain non-degeneracy conditions, the system converges to a unique invariant measure on (0, ∞)^*n*^. This condition for stochastic persistence is equivalent to the existence of positive weights *p*_1_, *p*_2_, …, *p*_*n*_ associated such that ∑_*i*_ *p*_*i*_*r*_*i*_(*μ*) *>* 0 for every stationary distribution *μ*(*dx*) on the extinct set. This latter formulation is known as the Hofbauer criterion for coexistence and was initially developed for systems of differential equations (Hofbauer, 1981). It was extended to stochastic difference and differential equations on compact domains first in (Schreiber et al., 2011) and extended to non-compact domains by (Hening and Nguyen, 2018*c*).

When a subset *I* ⊂ {1, …, *n*} of species coexist a, the long-term per-capita growth rate of each species in *I* is zero i.e.

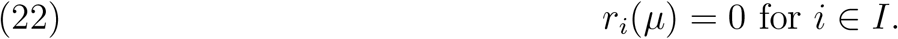

This implies that the mean species densities at stationarity, 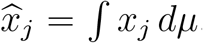, satisfy the following linear system of equations

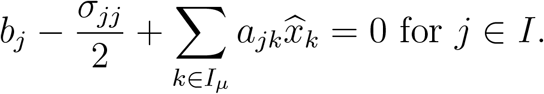

Solving the system, one can then obtain the invasion rates

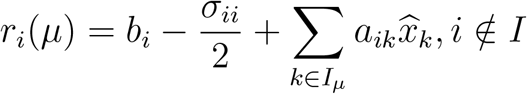

In particular, if *μ* is an invariant measure on (0, ∞)^*n*^, we have *r*_*i*_(*μ*) = 0 for any *i ∈* {1,, *n*}, which leads to the system

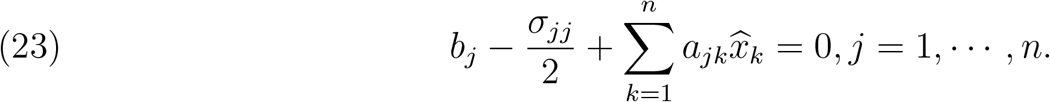

In the specific setting from (3), and remembering the identities from (21), if we set 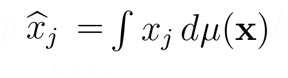 and use equation (23) we get that the average densities have to satisfy

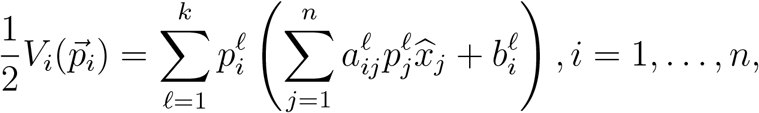

which is equivalent to (5).

## Appendix B. Characterizing the coESS

Coevolutionary Stable Strategies. To characterize a coESS of patch-selection, we consider the situation where mutations arise in a subset, *M* ⊂ {1, 2, …, *n*}, of the species. For these mutant species *i* ∈ *M*, let 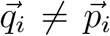 denote their patch-selection strategy and *y*_*i*_ their total population density. The dynamics of this augmented community are given by

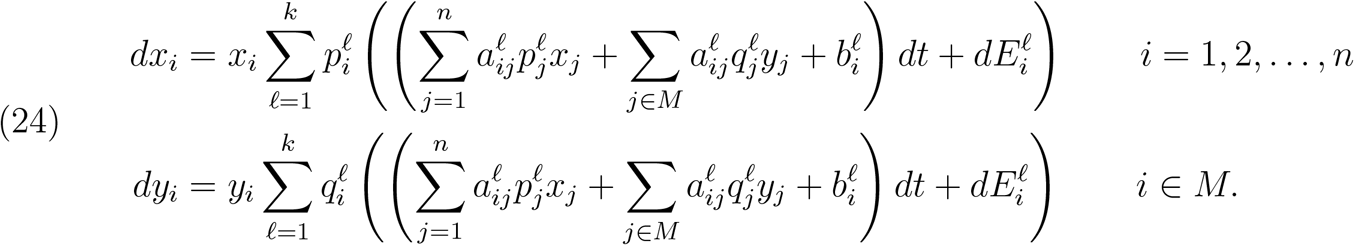

and let 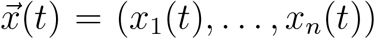 and 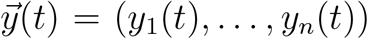. For an *n* × *n* symmetric matrix *Q*, define (note that the supremum is taken over nonpositive vectors)

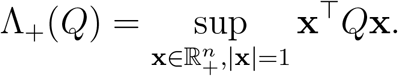

### Assumption B.1.

*(Boundedness) For any* 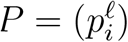, *let A*_*P*_ = (*a*_*ij*_)_*n*×*n*_ *where*

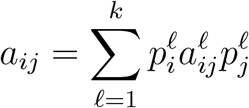

*There exists a diagonal matrix* 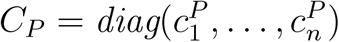 *with positive diagonal entries such that*

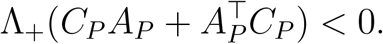

The next result shows that the boundedness assumption is satisfied for the examples we consider in the main text.

### Proposition B.1.

*Assumption B.1 holds for the predator-prey model* (31) *and for the competitive model* (16).

*Proof*. In the predator-prey model (31) we have

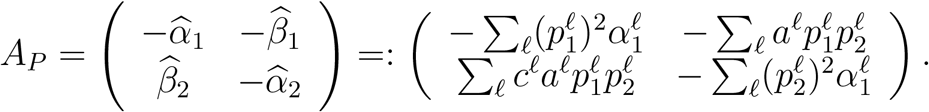

If we set 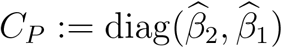 then

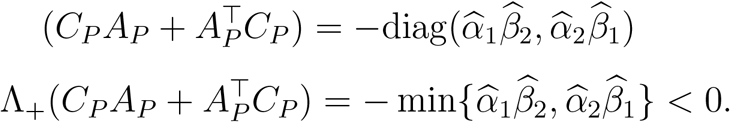

Similarly, since all the interaction coefficients of the competition model (16) are negative, by setting *C*_*P*_ = diag(1, 1) one can show that

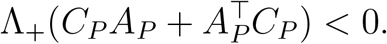

□

### Assumption B.2.

*(Persistence) For any invariant probability measure ν on* 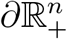 *we have*

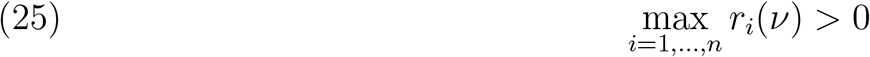

*where*

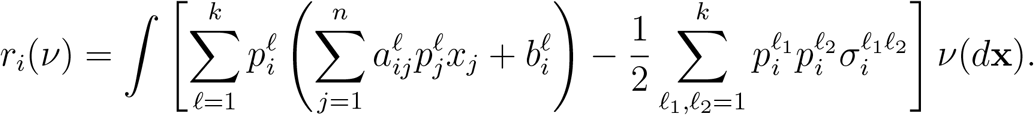

If *μ* is an invariant probability measure, define

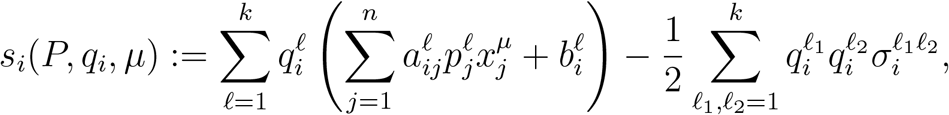

where 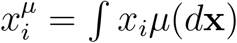. Note that the 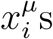 will be solutions to (5).

### Assumption B.3.

*(coESS) For any ergodic invariant probability measure μ of* (3) *on* 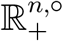 *we have*

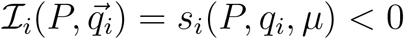

*for any q*_*i*_ ≠ *p*_*i*_.

### Theorem B.1.

*Suppose the strategy P satisfies Assumptions B.1 and B.2. Then the system* (3) *is persistent. If, in addition, the strategy P satisfies Assumption B.3, then for any set M* ⊂ {1, …, *n*} *of mutants with* 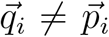 *for i* ∈ *M and ε >* 0, *there exists δ >* 0 *such that if y*_*i*_(0) ≤ *δ for all i, y*_*i*_(0) = 0 *for i with p*_*i*_ = *q*_*i*_, *and x*_*i*_(0) *>* 0 *for all i then with probability greater than* 1 − *ε the mutants go asymptotically extinct, i.e. y*_*m*_(*t*) → 0 *as t* → ∞, *and the residents persist, i.e. x*_*i*_(*t*) ↛ 0 *as t* → ∞ *for all i*.

*Proof*. Consider the system (24). Define

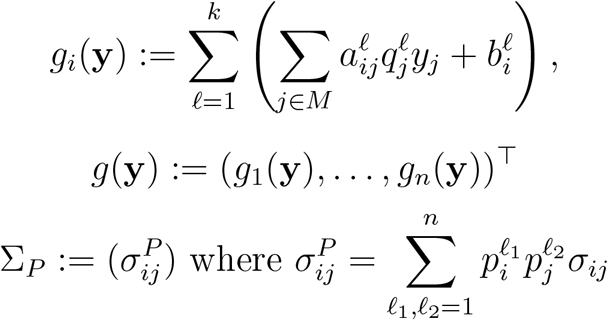

and

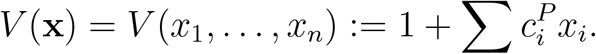

If ℒ is the generator of the process 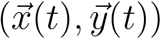 we have

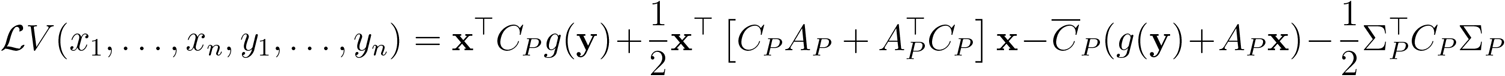

where 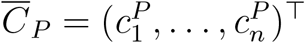. Using Assumption B.1

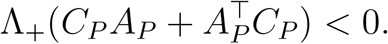

As a result we can find *K*_1_, *K*_2_ *>* 0 such that

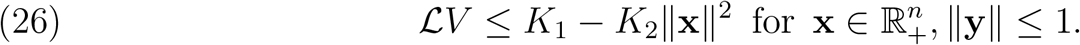

When **y** = 0, (26) implies that (Hening and Nguyen, 2018*a*, Assumption 1.1) holds. Note that Assumption B.2 is exactly (Hening and Nguyen, 2018*a*, Assumption 1.2). As a result, we can apply (Hening and Nguyen, 2018*a*, Theorem 4.1) in order to get that (3) is persistent.

Next, suppose that Assumption B.3 also holds. Under Assumptions B.2 and B.3, and the arguments from the begining of Section 5 in Hening and Nguyen (2018*a*), we can find sufficiently small constants 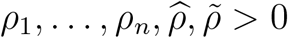 such that

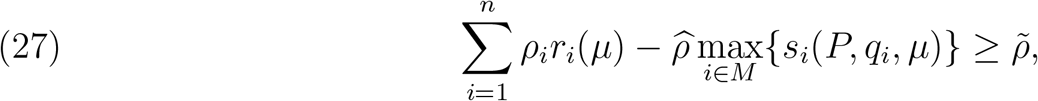

for any invariant measure *μ* of 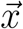 on 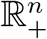. Consider the function

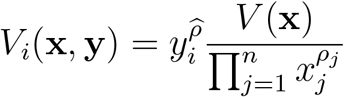

Because of (26), we can show that there is *H >* 0 such that

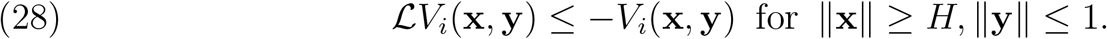

In view of (27) an d(28) and the arguments from Proposition 5.1 and Theorem 5.1 in Hening and Nguyen (2018*a*), we can show that there exist *δ >* 0, *T >* 0, *θ >* 0 such that for any starting point (**x, y**) 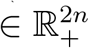

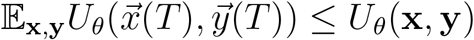

where

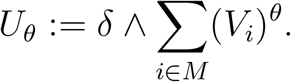

The proof can be finished by proceeding in the same manner as in (Hening and Nguyen, 2018*a*, Theorem 5.1 and Lemma 5.9). □

### Proposition B.2.

*Suppose the covariance matrices* 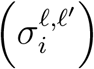, *are non-degenerate. If there exists* 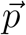*such that* 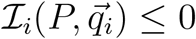 *for all i and* 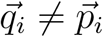 *then P is a co-ESS*.

*Proof*. Once we fix *p* the quadratic maximization problem 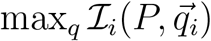 under the constraint ∑*q*_*i*_ = 1 has a unique maximum because the covariance matrix is positive definite. Since, by assumption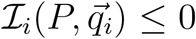, and 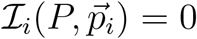 the unique maximum is at 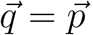. This implies that 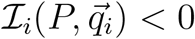 if 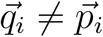. Therefore, *P* is a co-ESS. □

## Appendix C.

As getting an explicit tractable form of the coESS is difficult, we introduce a coevolutionary dynamic that can be used to solve for the coESS numerically. The basic idea behind the coevolutionary dynamic is that mutations arise in each species. These mutations randomly reallocate time spent in one patch with time spent in another patch. This reassignment is only “adopted” by species *i* if it increases its invasion growth rate 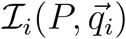. More formally, if time is discretized into units of length *ϵ* > 0 and *e*^*ℓ*^ denotes the standard unit vector whose *ℓ*-th component is 1 and remaining components are 0, then a mutation from 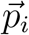 to 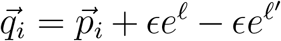 occurs with probability 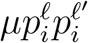 and establishes with probability

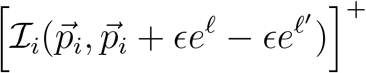

where *x*^+^ = max{0, *x*} denotes the positive part of a real number. The establishment probability is consistent with standard branching process approximations. The mutation probability is a choice of convenience. More general forms of mutation probabilities can be used. More general scalings proportional to *ϵ* simply correspond to rescaling time. Under these assumptions,

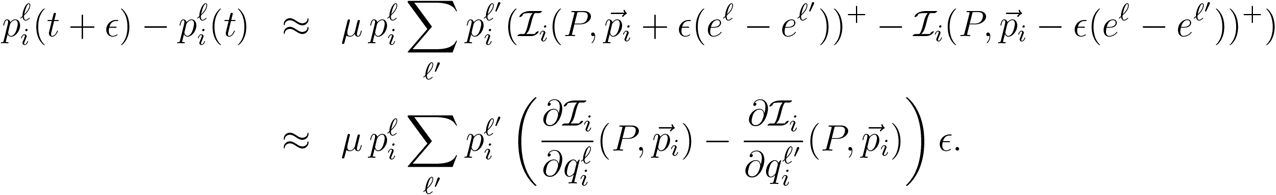

Hence, dividng by *ϵ* and taking the limit as *ϵ* → 0 yields

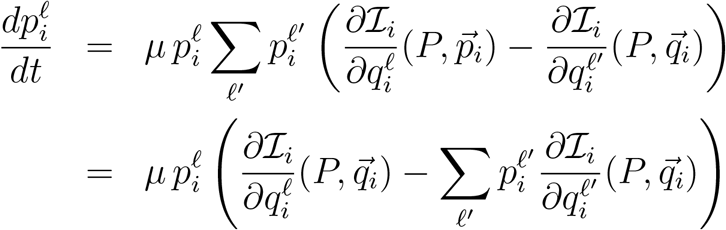

as claimed in the main text. Moreover, as

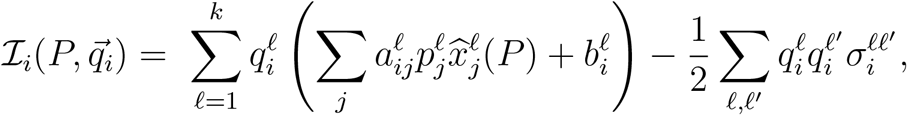

we have that

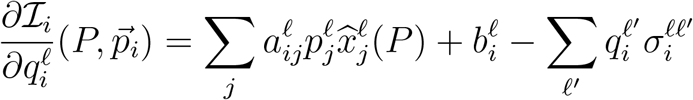

which provides explicit expressions for the coevolutionary dynamics.

We claim that if 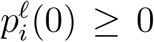 for all *ℓ* and 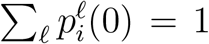, then 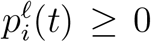 for all *ℓ* with *t* ≥ 0 and 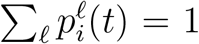 for all *t* ≥ 0. This claim follows from two observations. First, the system of equations is conservative as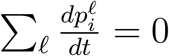. Hence, if 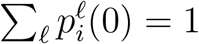, then 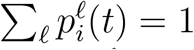 for all *t* ≥ 0. Second, since 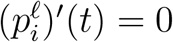 whenever 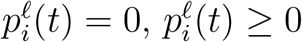 for *t* provided that 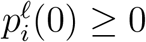.

Finally, consider any equilibrium 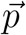 of the coevolutionary dynamic. If we have a species *i* such that 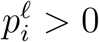 for all *ℓ*, then

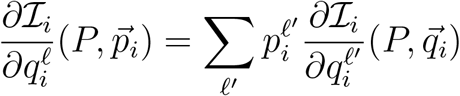

for all *ℓ*. Moreover,

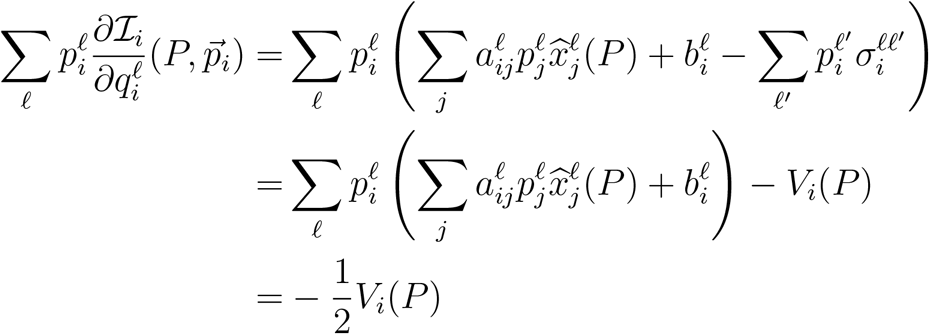

where the final inequality follows from the fact that 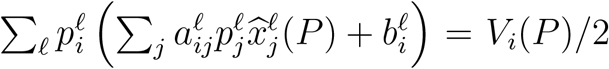 at the stationary distribution.

## Appendix D.

To derive a necessary condition for a co-ESS, we can use the method of Lagrange multipliers. Due to the constraint 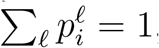, any co-ESS 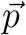 must satisfy

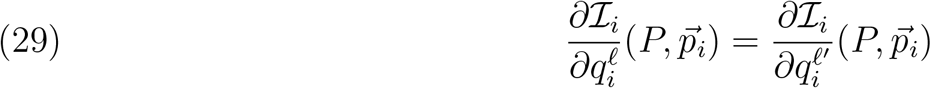

for any *i* and pair of occupied patches *ℓ, ℓ*′ by species *i* i.e.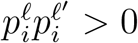. Computing these derivatives we get

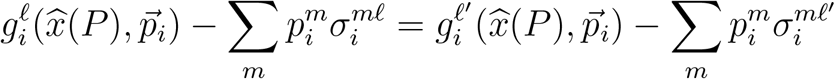

when 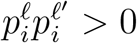. Now, if we multiply everything by 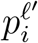and sum over *ℓ*′ we get, using equation (5), that the coESS must satisfy

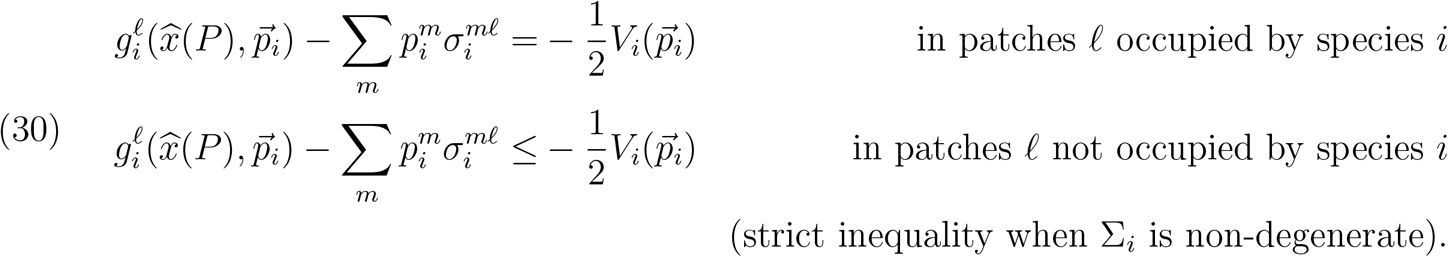

### Proposition D.1.

*If the covariance matrix for species i is non-degenerate, then the local stochastic growth rates (at the coESS) are negative in all patches *ℓ* occupied by species i, as long as more than one patch is occupied:* 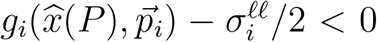 *when* 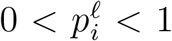. *If* 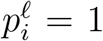 *then* 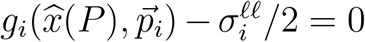. *As a result, all occupied patches are sinks or pseudo-sinks for species i*.

*Proof*. Combine Lemma D.1 below with the first equation from (9). □

### Lemma D.1.

*Let* 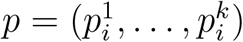 *be a vector in the k dimensional simplex* Δ, *so* 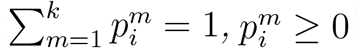. *Let ℓ* ∈ {1, …, *k*}. *Define*

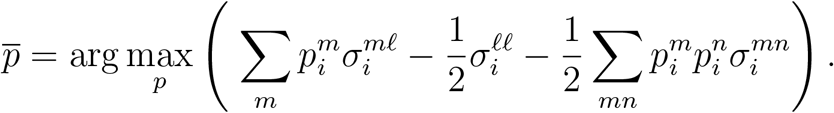

*Then* 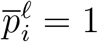, *and* 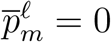 *for m* ≠ *ℓ*.

*Proof*. Since constant terms do not change where the maximum of an expression will be, we will drop the term 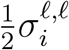 and look at

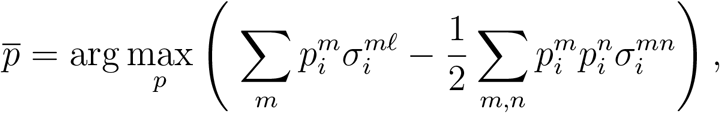

Since ∑ is positive definite we can write it as ∑ = Γ^*t*^Γ for Γ invertible. Let *X* be the unit vector in the *ℓ*–th direction i.e. *X*_*ℓ*_ = 1 and *X*_*m*_ = 0 for *m* ≠ *ℓ*. Then the maximization problem can be written as:

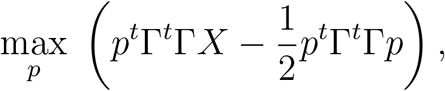

subject to *p* ∈ Δ. Now simplify the expression by defining *Y* := Γ*p*, and *L* := Γ*X*. Since Γ is invertible we rewrite the maximization problem as

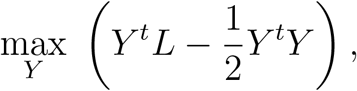

subject to the Γ^−1^*Y* ∈ Δ. Finding the maximizing *Y* is equivalent to finding the maximizing *p* in the previous expression.

Let *Ŷ* = *Y/║Y ║* be the unit vector in the direction of *Y* and *y* = ║*Y ║*. The maximization problem becomes

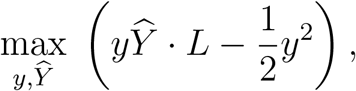

subject to Γ^−1^*Y* ∈ Δ. It is clear that *y >* 0. We can therefore see that, given any fixed *y >* 0, the maximum is when *Ŷ* is parallel to *L*. This implies that 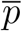 and *X* are parallel, and from the sum condition, we deduce that in fact 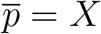. □

### Proposition D.2.

*Only one patch is occupied, say patch *ℓ*, by species i at the coESS if and only if*

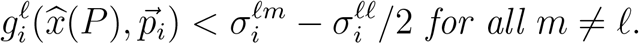

*Proof*. Suppose that 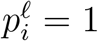for the coESS for species *i*. Consider any probability vector 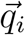 such that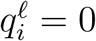. Define

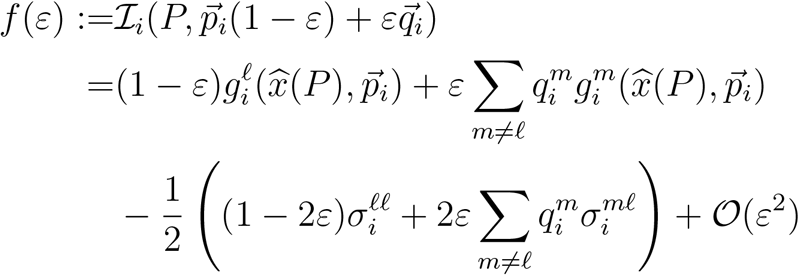

Taking the derivative with respect to *ε* and evaluating at *ε* = 0, the coESS must satisfy

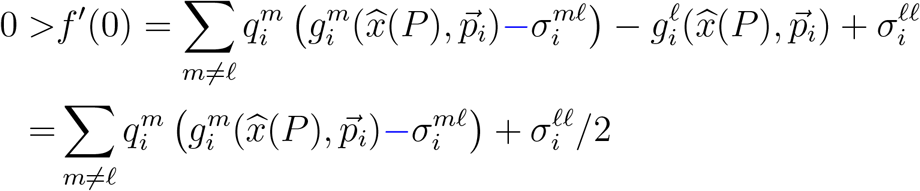

where the second line follows from the local stochastic growth rate 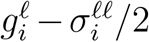 of species *i* equally zero in patch *ℓ* if that is the only patch being occupied. By linearity, this inequality holds for all choices of 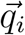 with 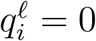 if and only if it holds for all 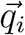 satisfying 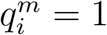 for *m* ≠ *ℓ*. □

### Proposition D.3.

*If the covariance matrix is a positive diagonal matrix for species i, then* 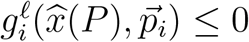 *for any patch *ℓ* not occupied by species i (i.e*.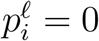*). In particular, this implies that there are no unoccupied stochastic sinks for species i, i.e. patches where* 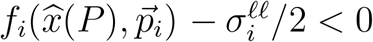 *and* 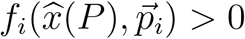.

*Proof*. We want to maximize

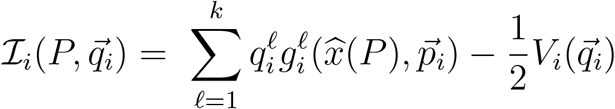

with the constraint 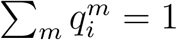 and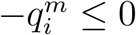. Let 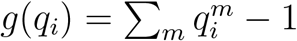 and 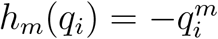.

We know that the maximum is at 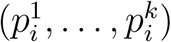. It has to satisfy the conditions

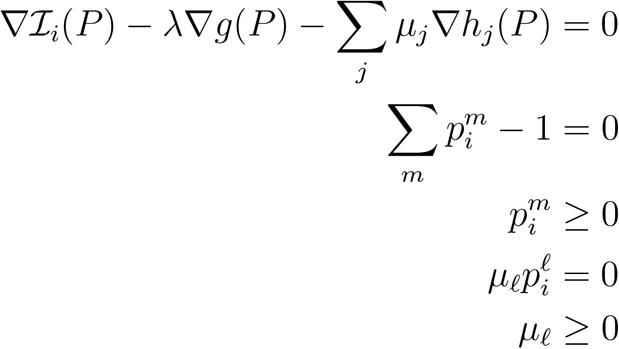

If 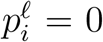 and using that ∑ is diagonal together with (29) and (9) we get that 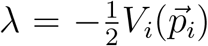 and

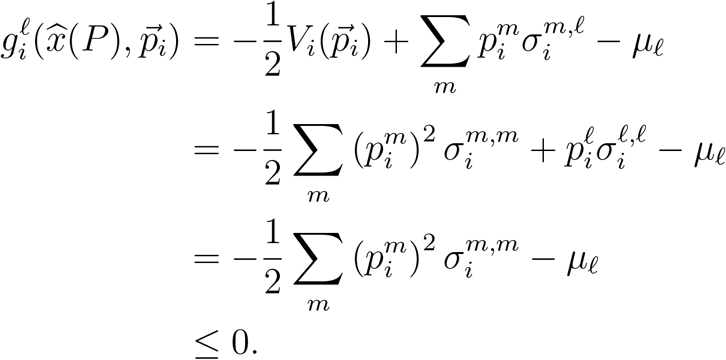

□

## Appendix E.

The general model for predator-prey interactions considers a predator with global density *x*_2_ and its prey with global density *x*_1_. In each patch, the prey exhibit an intrinsic rate of growth *r*^*ℓ*^ and intraspecific competition coefficient 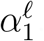. The predator, in patch *ℓ*, attacks the prey with an attack rate *a*^*ℓ*^, experiences a per-capita death rate *d*^*ℓ*^, and an intraspecific competition coefficient 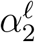. Prey captured in patch *ℓ* are converted with efficiency *c*^*ℓ*^ to predator offspring.

The predator-prey dynamics become

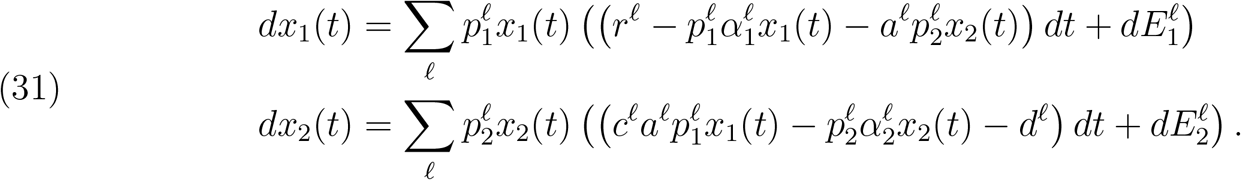

The predator-prey model persists if

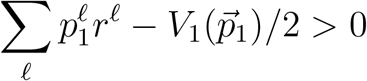

and

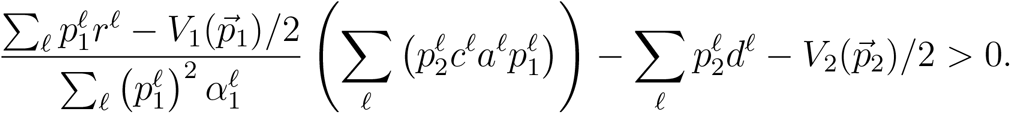

This result follows from Appendix A or from Hening and Nguyen (2018*c*). At stationarity

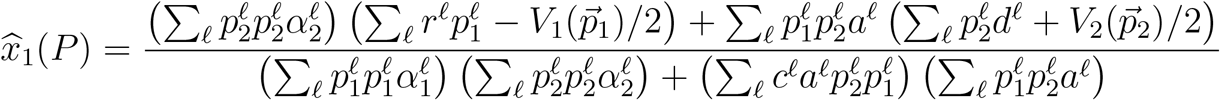

and

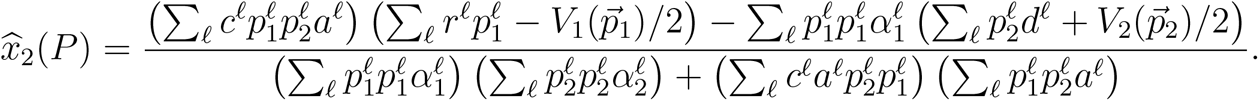

The invasion rate for a mutant prey is

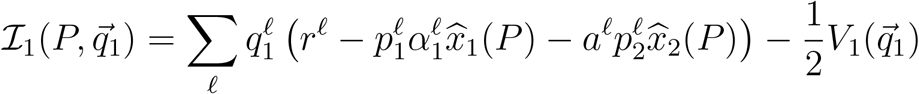

while the invasion rate for a mutant predator becomes

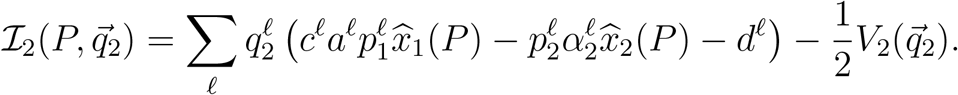

The strategy *P* is a coESS if for all *q* ≠ *P*

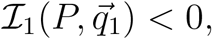

and

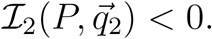

By the criterion from (30), a necessary condition for *p* to be a coESS is that for *i* = 1, 2

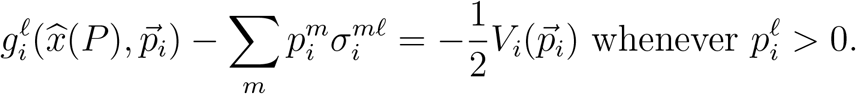

This means that

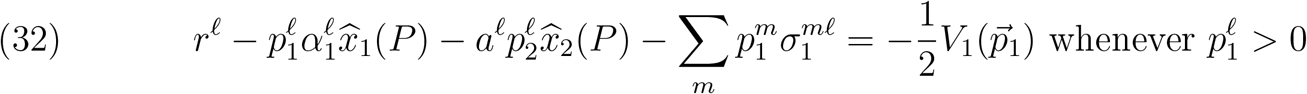

and

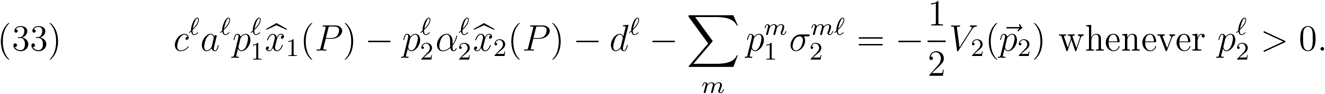

We next look at some specific examples.

### Example 1.

Suppose 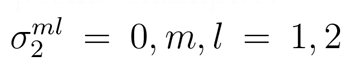 and 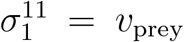 and 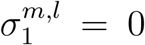 if (*m, l*) ≠ (1, 1) 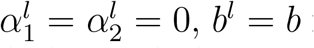 for *b* ∈ {*a, c, d*}, *l* = 1, 2, *r*^1^ = *r*_source_ *>* 0 and *r*^2^ = −*r*_sink_ *<* 0. In this setting (32) and (33) become

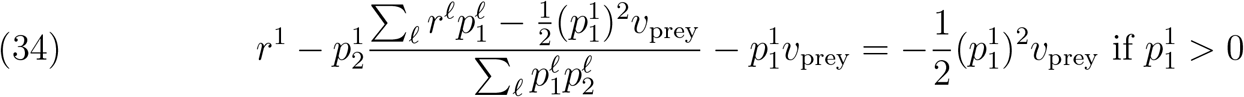

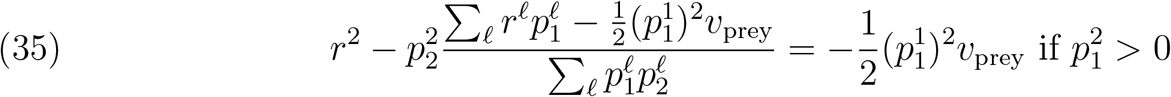

and

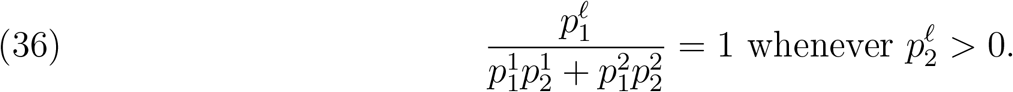

If *v*_prey_ *<* 2*r*_sink_ = 2*r*^2^ then (35) implies that 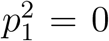 which, combined with (36), implies that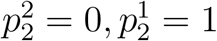.

If *v*_prey_ *<* 2*r* = 2*r*^2^ and 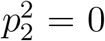 then (35) leads to 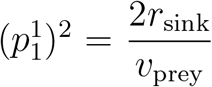. This formula holds true until

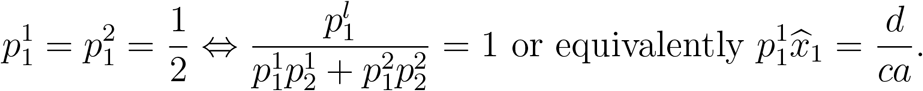

When 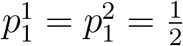, (34) and (35) become

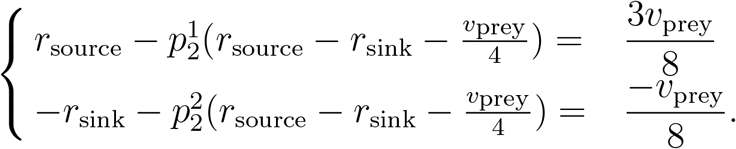

The above system has solutions satisfying 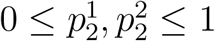 and 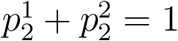 if and only if

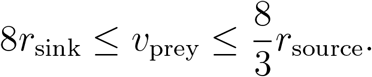

If 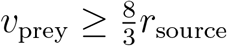 then (34) implies that 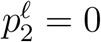. Using (34) again we get that

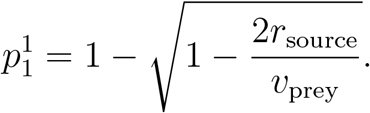

Meanwhile, 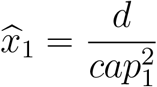 decreases as *v*_prey_ increases.

### Example 2.

When 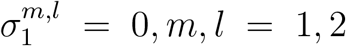 and 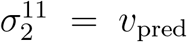 and 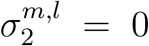 if (*m, l*) ≠ (1, 1) 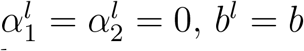 for *b* ∈ {*a, c, d*}, *l* = 1, 2, *r*^1^ = *r*_source_ *>* 0 and *r*^2^ = −*r*_sink_ *<* 0, (32) and (33) become

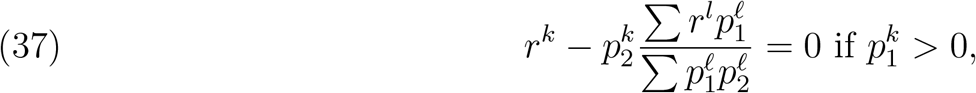

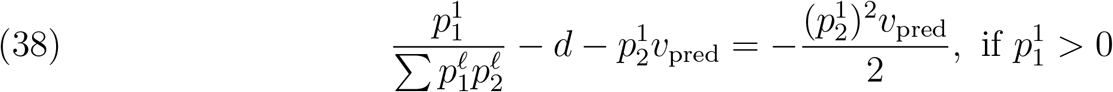

and

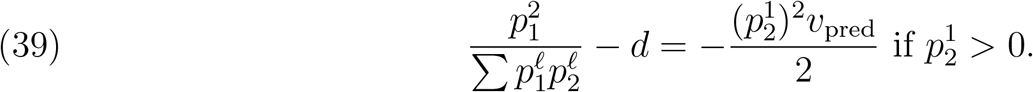

Since *r*^2^ = −*r*_sink_ *<* 0, it follows from (37) that 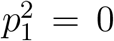. Then, 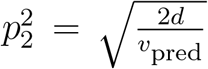 due to (39) if *v*_pred_ ≥ 2*d*.

## Appendix F.

The two species coexist if the following two conditions hold

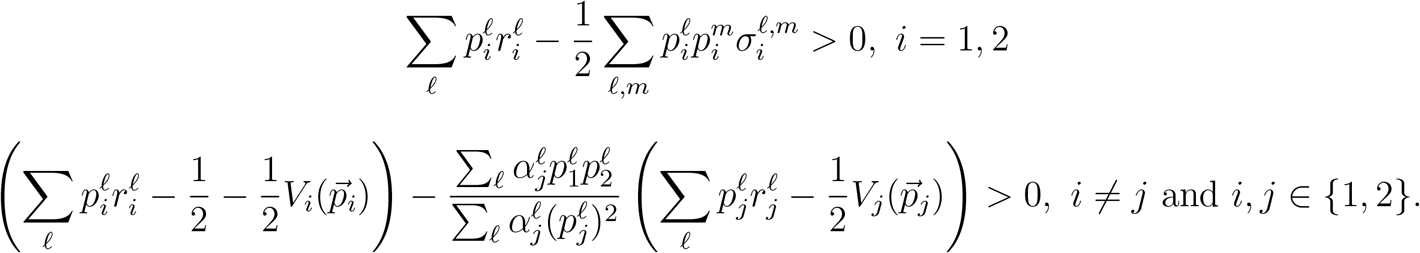

At stationarity we get

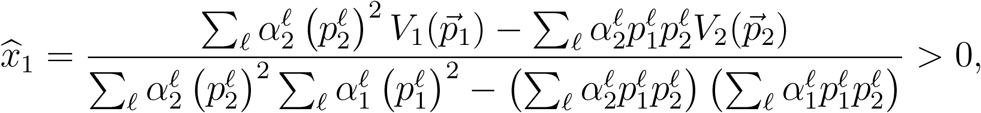

and

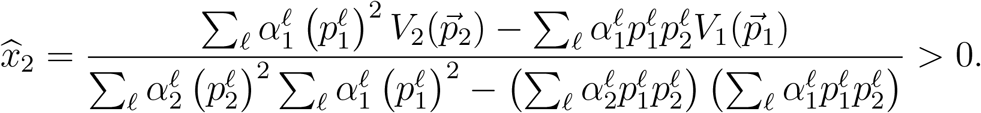

A necessary condition for a coESS is that for all strategies *q ≠ p*

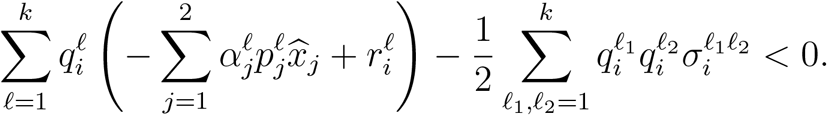

We get that a necessary condition for *p* to be a coESS is that for *i* = 1, 2

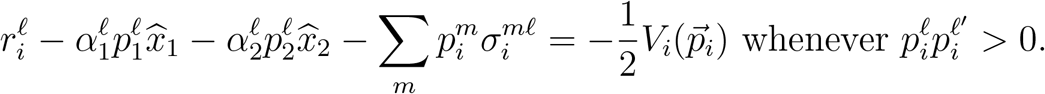

Now we analyze the symmetric case discussed in the main text. In this case, there is an even number of patches *k* = 2*m* for some *m*. As stated in the main text, the per-capita growth rate for the species with competitive advantage 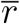 is greater than the per-capita growth rate <u>*r*</u> of the species with competitive disadvantage, 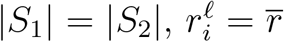 for *ℓ* ∉ *S*_*i*_, and 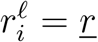 for *ℓ* ∉ *S*_*i*_. We also assume that the covariance matrix for both species is ∑_*i*_ = *vI* for *i* = 1, 2.

By symmetry, any coESS must satisfy 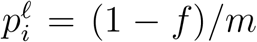 for *ℓ* ∈ *S*_*i*_ and *f/m* for *ℓ* ∉ *S*_*i*_. For any choice of *f*, let us first solve for the mean density of competitors within a patch 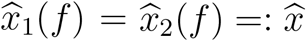 in which case the global mean density for species *i* is 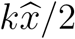. This mean density must satisfy

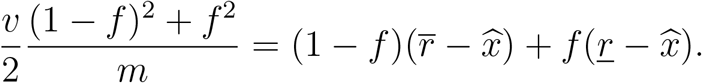

This implies

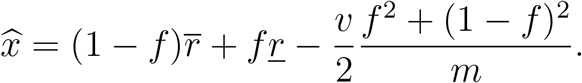

The invasion rate of a different strategy 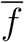 invading a resident community with strategy *f* equals

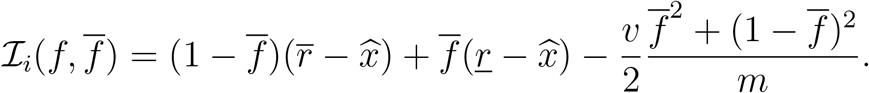

Hence,

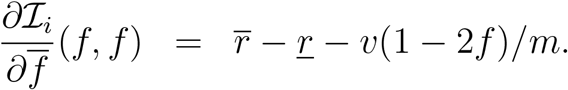

There are two cases to consider. Either *f* = 0 or 0 *< f <* 1. The first case only occurs if the partial derivative of ℐ_*i*_ at *f* = 0 is positive. This occurs if

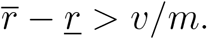

Hence, if the noise is too small, or there are many patches, or the difference in the competitive advantages are too large, then the species remain segregated at the coESS.

Consider the case when 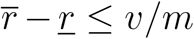. Then the coESS must satisfy that the partial derivative of ℐ_*i*_ at *f* = 0 equals zero. In this case, the coESS must satisfy

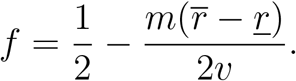

This equation implies that the majority of the individuals are in the patches in which they are competitively superior. However, for smaller differences in competitive ability and higher levels of stochasticity, individuals spread more equally across the patches.

## References

Abrams, P. A. 2009. When does greater mortality increase population size? the long history and diverse mechanisms underlying the hydra effect. Ecology Letters 12:462–474.

Amarasekare, P., and R. M. Nisbet. 2001. Spatial heterogeneity, source-sink dynamics, and the local coexistence of competing species. American Naturalist 158:572–584.

Barson, N., J. Cable, and C. Van Oosterhout. 2009. Population genetic analysis of microsatellite variation of guppies (poecilia reticulata) in trinidad and tobago: evidence for a dynamic source–sink metapopulation structure, founder events and population bottlenecks. Journal of evolutionary biology 22:485–497.

Bascompte, J., H. Possingham, and J. Roughgarden. 2002. Patchy populations in stochastic environments: Critical number of patches for persistence. American Naturalist 159:128—137.

Benaïm, M. 2018. Stochastic persistence. arXiv preprint arXiv:1806.08450.

Benaïm, M., and C. Lobry. 2016. Lotka–Volterra with randomly fluctuating environments or “how switching between beneficial environments can make survival harder”. The Annals of Applied Probability 26:3754–3785.

Benaïm, M., and S. Schreiber. 2019. Persistence and extinction for stochastic ecological models with internal and external variables. Journal of Mathematical Biology 79:393–431.

Berdegue, M., J. Trumble, J. Hare, and R. Redak. 1996. Is it enemy-free space? the evidence for terrestrial insects and freshwater arthropods. Ecological Entomology 21:203–217.

Brown, J., and T. Vincent. 1987. Coevolution as an evolutionary game. Evolution 41:66–79.

Burner, R., A. Styring, M. Rahman, and F. Sheldon. 2019. Occupancy patterns and upper range limits of lowland bornean birds along an elevational gradient. Journal of Biogeography 46:2583–2596.

Campos-Cerqueira, M., W. Arendt, J. Wunderle Jr, and T. Aide. 2017. Have bird distributions shifted along an elevational gradient on a tropical mountain? Ecology and Evolution 7:9914– 9924.

Cantrell, R., and C. Cosner. 2018. Evolutionary stability of ideal free dispersal under spatial heterogeneity and time periodicity. Mathematical Biosciences 305:71–76.

Cantrell, R., C. Cosner, and K. Lam. 2021. Ideal free dispersal under general spatial hetero-geneity and time periodicity. SIAM Journal on Applied Mathematics 81:789–813.

Chesson, P., and J. Kuang. 2008. The interaction between predation and competition. Nature 456:235–238.

Chesson, P. L. 1982. The stabilizing effect of a random environment. Journal of Mathematical Biology 15:1–36.

Chettri, S., B.and Bhupathy, and B. Acharya. 2010. Distribution pattern of reptiles along an eastern himalayan elevation gradient, india. Acta Oecologica 36:16–22.

Childs, D., C. Metcalf, and M. Rees. 2010. Evolutionary bet-hedging in the real world: empirical evidence and challenges revealed by plants. Proceedings of the Royal Society B: Biological Sciences 277:3055–3064.

Cohen, D. 1966. Optimizing reproduction in a randomly varying environment. J. Theoretical Biology 12:119–129.

Cole, N., C. Jones, and S. Harris. 2005. The need for enemy-free space: the impact of an invasive gecko on island endemics. Biological Conservation 125:467–474.

Connell, J. 1980. Diversity and the coevolution of competitors, or the ghost of competition past. Oikos pages 131–138.

Cortez, M. H., and P. A. Abrams. 2016. Hydra effects in stable communities and their implications for system dynamics. Ecology 97:1135–1145.

Denno, R., S. Larsson, and K. Olmstead. 1990. Role of enemy-free space and plant quality in host-plant selection by willow beetles. Ecology 71:124–137.

Diamond, J. 1973. Distributional ecology of new guinea birds: recent ecological and biogeographical theories can be tested on the bird communities of new guinea. Science 179:759–769.

Diamond, J. 1978. Niche shifts and the rediscovery of interspecific competition: Why did field biologists so long overlook the widespread evidence for interspecific competition that had already impressed darwin? American scientist 66:322–331.

Dias, P., G. Verheyen, and M. Raymond. 1996. Source-sink populations in Mediterranean Blue tits: evidence using single-locus minisatellite probes. Journal of Evolutionary Biology 9:965– 978.

Edwards, K. F., and S. J. Schreiber. 2010. Preemption of space can lead to intransitive coexistence of competitors. Oikos 119:1201–1209.

Evans, S., A. Hening, and S. J. Schreiber. 2015. Protected polymorphisms and evolutionary stability of patch-selection strategies in stochastic environments. Journal of Mathematical Biology 71:325–359.

Evans, S. N., P. Ralph, S. J. Schreiber, and A. Sen. 2013. Stochastic growth rates in spatiotemporal heterogeneous environments. Journal of Mathematical Biology 66:423–476.

Feng, T., C. Li, X. Zheng, S. Lessard, and Y. Tao. 2022. Stochastic replicator dynamics and evolutionary stability. Physical Review E 105:044403.

Foster, D., and P. Young. 1990. Stochastic evolutionary game dynamics. Theoretical population biology 38:219–232.

Fox, L. R., and J. Eisenbach. 1992. Contrary choices: possible exploitation of enemy-free space by herbivorous insects in cultivates vs. wild crucifers. Oecologia 89:574–579.

Fretwell, S. D., and H. L. J. Lucas. 1969. On territorial behavior and other factors influencing habitat distribution in birds. Acta Biotheoretica 19:16–36.

Furrer, R., and G. Pasinelli. 2016. Empirical evidence for source–sink populations: a review on occurrence, assessments and implications. Biological Reviews 91:782–795.

Gardiner, C. 2009. Stochastic methods: A handbook for the natural and social sciences. Series in Synergetics, 4th ed. Springer, Berlin.

Greeney, H., M. Meneses, C. Hamilton, E. Lichter-Marck, R. W. Mannan, N. Snyder, H. Snyder, S. Wethington, and L. Dyer. 2015. Trait-mediated trophic cascade creates enemy-free space for nesting hummingbirds. Science advances 1:e1500310.

Hänfling, B., and D. Weetman. 2006. Concordant genetic estimators of migration reveal anthropogenically-enhanced source-sink population structure in the river sculpin, Cottus gobio. Genetics 173:1487–1501.

Hassell, M. P., R. M. May, S. W. Pacala, and P. L. Chesson. 1991. The persistence of host-parasitoid associations in patchy environments. I. A general criterion. American Naturalist 138:586–583.

Heisswolf, A., E. Obermaier, and H. Poethke. 2005. Selection of large host plants for oviposition by a monophagous leaf beetle: nutritional quality or enemy-free space? Ecological Entomology 30:299–306.

Hening, A., and D. H. Nguyen. 2018a. Coexistence and extinction for stochastic Kolmogorov systems. Annals of Applied Probability 28:1893–1942.

Hening, A., and D. H. Nguyen. 2018b. Persistence in stochastic Lotka-Volterra food chains with intraspecific competition. Bulletin of Mathematical Biology 80:2527–2560.

Hening, A., and D. H. Nguyen. 2018c. Stochastic Lotka–Volterra food chains. Journal of Mathematical Biology 77:135– 163.

Hening, A., D. H. Nguyen, and P. Chesson. 2021. A general theory of coexistence and extinction for stochastic ecological communities. Journal of Mathematical Biology 82:1–76.

Hofbauer, J. 1981. A general cooperation theorem for hypercycles. Monatshefte für Mathematik 91:233–240.

Hofbauer, J., and K. Sigmund. 1998. Evolutionary Games and Population Dynamics. Cambridge University Press.

Holt, R. 1977. Predation, apparent competition and the structure of prey communities. Theoretical Population Biology 12:197–229.

Holt, R. 1985. Patch dynamics in two-patch environments: Some anomalous consequences of an optimal habitat distribtuion. Theoretical Population Biology 28:181–208.

Holt, R. D. 1997. On the evolutionary stability of sink populations. Evolutionary Ecology 11:723–731.

Hutson, V., K. Mischaikow, and P. Poláčik. 2001. The evolution of dispersal rates in a heterogeneous time-periodic environment. Journal of Mathematical Biology 43:501–533.

Jansen, V. A. A., and J. Yoshimura. 1998. Populations can persist in an environment consisting of sink habitats only. Proceeding of the National Academy of Sciences USA 95:3696–3698.

Jeffries, M. J., and J. H. Lawton. 1984. Enemy-free space and the structure of ecological communities. Biological Journal of the Linnean Society 23:269–86.

Jonzén, N., C. Wilcox, and H. Possingham. 2004. Habitat selection and population regulation in temporally fluctuating environments. American Naturalist 164:103–103.

Kadmon, R., and K. Tielbörger. 1999. Testing for source-sink population dynamics: an experimental approach exemplified with desert annuals. Oikos 86:417–429.

Kaminski, L., A. Freitas, and P. Oliveira. 2010. Interaction between mutualisms: ant-tended butterflies exploit enemy-free space provided by ant-treehopper associations. The American Naturalist 176:322–334.

Keagy, J., S. J. Schreiber, and D. A. Cristol. 2005. Replacing sources with sinks: When do populations go down the drain? Restoration Ecology 13:529–535.

Kisdi, E. 2002. Dispersal: Risk spreading versus local adaptation. The American Naturalist 159:579–596.

Kreuzer, M., and N. Huntly. 2003. Habitat-specific demography: evidence for source-sink population structure in a mammal, the pika. Oecologia 134:343–349.

Křivan, V., R. Cressman, and C. Schneider. 2008. The ideal free distribution: a review and synthesis of the game-theoretic perspective. Theoretical Population Biology 73:403–425.

Lande, R., S. Engen, and B. Sæther. 2003. Stochastic population dynamics in ecology and conservation: An introductions. Oxford University Press.

Law, R., and R. D. Morton. 1996. Permanence and the assembly of ecological communities. Ecology 77:762–775.

Lawlor, L., and J. Maynard Smith. 1976. The coevolution and stability of competing species. The American Naturalist 110:79–99.

Levin, S. A., D. Cohen, and A. Hastings. 1984. Dispersal strategies in patchy environments. Theoretical Population Biology 26:165 – 191.

Lion, S., and S. Gandon. 2015. Evolution of spatially structured host–parasite interactions. Journal of Evolutionary Biology 28:10–28.

Lion, S., M. Van Baalen, and W. Wilson. 2006. The evolution of parasite manipulation of host dispersal. Proceedings of the Royal Society B: Biological Sciences 273:1063–1071.

Loreau, M., T. Daufresne, A. Gonzalez, D. Gravel, F. Guichard, S. Leroux, N. Loeuille, F. Massol, and N. Mouquet. 2013. Unifying sources and sinks in ecology and e arth sciences. Biological Reviews 88:365–379.

Manier, M., and S. Arnold. 2005. Population genetic analysis identifies source–sink dynamics for two sympatric garter snake species (Thamnophis elegans and Thamnophis sirtalis). Molecular Ecology 14:3965–3976.

Markowitz, H. 1952. Portfolio selection. The Journal of Finance 7:77–91.

Markowitz, H. M. 1991. Foundations of portfolio theory. The Journal of Finance 46:469–477.

Markowitz, H. M. 2010. Portfolio theory: As i still see it. Annual Review of Financial Economics 2:1–23.

May, R. M. 1973. Stability in randomly fluctuating versus deterministic environments. The American Naturalist 107:621–650.

May, R. M. 1975. Stability and Complexity in Model Ecosystems, 2nd edn. Princeton University Press, Princeton.

McDowall, R. 2010. Why be amphidromous: expatrial dispersal and the place of source and sink population dynamics? Reviews in Fish Biology and Fisheries 20:87–100.

McPeek, M., and R. D. Holt. 1992. The evolution of dispersal in spatially and temporally varying environments. American Naturalist 6:1010–1027.

Monson, D., D. Doak, B. Ballachey, and J. Bodkin. 2011. Could residual oil from the exxon valdez spill create a long-term population “sink” for sea otters in alaska? Ecological Applications 21:2917–2932.

Morris, D. 2011. Adaptation and habitat selection in the eco-evolutionary process. Proceedings of the Royal Society B: Biological Sciences 278:2401–2411.

Morris, D. W. 2003. Toward an ecological synthesis: a case for habitat selection. Oecologia 136:1–13.

Murphy, S. 2004. Enemy-free space maintains swallowtail butterfly host shift. Proceedings of the National Academy of Sciences 101:18048–18052.

Murphy, S., J. Lill, M. Bowers, and M. Singer. 2014. Enemy-free space for parasitoids. Environmental entomology 43:1465–1474.

Nolting, B. C., and K. C. Abbott. 2016. Balls, cups, and quasi-potentials: quantifying stability in stochastic systems. Ecology 97:850–864.

Noon, B. 1981. The distribution of an avian guild along a temperate elevational gradient: the importance and expression of competition. Ecological Monographs 51:105–124.

Oksanen, T., M. Power, and L. Oksanen. 1995. Ideal free habitat selection and consumer-resource dynamics. American Naturalist 146:565–585.

Oksendal, B. 2013. Stochastic differential equations: an introduction with applications. Springer Science & Business Media.

Polis, G. A., and R. D. Holt. 1992. Intraguild predation: The dynamics of complex trophic interactions. Trends in Ecology and Evolution 7:151–154.

Pulliam, H., and B. Danielson. 1991. Sources, sinks and habitat selection: A landscape perspective on population dynamics. American Naturalist 137:S50–S66.

Pulliam, H. R. 1988. Sources, sinks, and population regulation. American Naturalist 132:652– 661.

Rand, D. A., H. B. Wilson, and J. M. McGlade. 1994. Dynamics and evolution: evolutionary stable attractors, invasion exponents and phenotype dynamics. Phil. Trans. R. Soc. Lond. B 343:261–283.

Robinson, H., R. Wielgus, H. Cooley, and S. Cooley. 2008. Sink populations in carnivore management: cougar demography and immigration in a hunted population. Ecological Applications 18:1028–1037.

Rohr, R., S. Saavedra, G. Peralta, C. M. Frost, L.-F. Bersier, J. Bascompte, and J. Tylianakis. 2016. Persist or produce: a community trade-off tuned by species evenness. The American Naturalist 188:411–422.

Ronce, O. 2007. How does it feel to be like a rolling stone? ten questions about dispersal evolution. Annual Review of Ecology and Systematics 38:231–253.

Rosenzweig, M. 1981. A theory of habitat selection. Ecology 62:327–335.

Rosenzweig, M. 1987. Habitat selection as a source of biological diversity. Evolutionary Ecology pages 315–330.

Rosenzweig, M. 1991. Habitat selection and population interactions: the search for mechanism. The American Naturalist pages S5–S28.

Roughgarden, J. 1979. Theory of population genetics and evolutionary ecology. Macmillan.

Rowe, C., W. Hopkins, and V. Coffman. 2001. Failed recruitment of southern toads (bufo terrestris) in a trace element-contaminated breeding habitat: direct and indirect effects that may lead to a local population sink. Archives of Environmental Contamination and Toxicology 40:399–405.

Roy, H., L. Handley, K. Schönrogge, R. Poland, and B. Purse. 2011. Can the enemy release hypothesis explain the success of invasive alien predators and parasitoids? BioControl 56:451– 468.

Rubinstein, M. 2002. Markowitz’s “portfolio selection”: A fifty-year retrospective. The Journal of Finance 57:1041–1045.

Schmidt, K., J. Earnhardt, J. Brown, and R. Holt. 2000. Habitat selection under temporal heterogeneity: Exorcizing the ghost of competition past. Ecology 81:2622–2630.

Schreiber, S. 2012. Evolution of patch selection in stochastic environments. American Naturalist 180:17–34.

Schreiber, S., L. Fox, and W. Getz. 2000. Coevolution of contrary choices in host-parasitoid systems. American Naturalist pages 637–648.

Schreiber, S., S. Patel, and C. terHorst. 2018. Evolution as a coexistence mechanism: Does genetic architecture matter? The American Naturalist 191:407–420.

Schreiber, S., and M. Vejdani. 2006. Handling time promotes the coevolution of aggregation in predator-prey systems. Proceedings of the Royal Society: Biological Sciences 273:185–191.

Schreiber, S. J., M. Benaïm, and K. A. S. Atchadé. 2011. Persistence in fluctuating environments. Journal of Mathematical Biology 62:655–683.

Schreiber, S. J., L. R. Fox, and W. M. Getz. 2002. Parasitoid sex allocation affects coevolution of patch selection in host-parasitoid systems. Evolutionary Ecology Research 4:701–718.

Schreiber, S. J., and E. Saltzman. 2009. Evolution of predator and prey movement into sink habitats. American Naturalist 174:68–81.

Sieber, M., and F. M. Hilker. 2012. The hydra effect in predator–prey models. Journal of mathematical biology 64:341–360.

Soetaert, K., T. Petzoldt, and R. Setzer. 2010. Solving differential equations in R: Package deSolve. Journal of Statistical Software 33:1–25.

Stearns, S. C. 2000. Daniel Bernoulli (1738): evolution and economics under risk. Journal of Biosciences 25:221–228.

Tittler, R., L. Fahrig, and M. Villard. 2006. Evidence of large-scale source-sink dynamics and long-distance dispersal among Wood Thrush populations. Ecology 87:3029–3036.

Turelli, M. 1977. Random environments and stochastic calculus. Theoretical population biology 12:140–178.

Turelli, M. 1978. A reexamination of stability in randomly varying versus deterministic environments with comments on the stochastic theory of limiting similarity. Theoretical Population Biology 13:244–267.

Turelli, M. 1986. Stochastic community theory: a partially guided tour. Pages 321–339 in Mathe-matical Ecology. Springer.

Turelli, M., and J. H. Gillespie. 1980. Conditions for the existence of stationary densities for some two-dimensional diffusion processes with applications in population biology. Theoretical population biology 17:167–189.

Urban, M. C., S. Y. Strauss, F. Pelletier, E. P. Palkovacs, M. A. Leibold, A. P. Hendry, L. De Meester, S. M. Carlson, A. L. Angert, and S. T. Giery. 2020. Evolutionary origins for ecological patterns in space. Proceedings of the National Academy of Sciences.

van Baalen, M., and M. W. Sabelis. 1993. Coevolution of patch selection strategies of predator and prey and the consequences for ecological stability. American Naturalist 142:646–670.

Vierling, K. T. 2000. Source and sink habitats of red-winged blackbirds in a rural/suburban landscape. Ecological Applications 10:1211–1218.

Watkinson, A., and W. Sutherland. 1995. Sources, sinks and pseudo-sinks. Journal of Animal Ecology 64:126–130.

Zivot, E. 2017. Introduction to computational finance and financial econometrics. Chapman & Hall Crc.

